# Discovery of an IKK2 Site that Allosterically Controls Its Activation

**DOI:** 10.1101/2021.01.27.428502

**Authors:** Sonjiala Jackson Hotchkiss, Maria Carmen Mulero, Garrett J. Chan, Tapan Biswas, Smarajit Polley, Christine Ohn, Srihari Konduri, Dionicio Siegel, Özlem Demir, Rommie E. Amaro, Gourisankar Ghosh

## Abstract

IκB kinase 2/β (IKK2) is a critical regulator of inflammation which is inducibly activated by a host of stimuli. Aberrant activation of IKK2 is the leading cause of most inflammatory diseases and many associated cancers. Efforts to prevent these diseases by small-molecule inhibitors of IKK2 activity have not been successful. Most inhibitors developed for IKK2 are ATP-competitive, and they are toxic *in vivo* due to their off-target effects. Here we focused on identifying inhibitors to block IKK2 activity from an allosteric site, not the ATP-binding pocket. Using virtual screening, we first identified several candidate allosteric sites and screened for potential small-molecule binders, and then selected candidates inhibitory to IKK2 activity using cell-based functional assays. Hydrogen deuterium exchange coupled to mass-spectrometry (HDX-MS) and MS-MS assays revealed that a class of benzoyl conjugates of pyrrolidinedione covalently bound to a site located at the interface of the kinase domain (KD) and the helical domain (SDD), and inhibited IKK2 activation allosterically by preventing phosphorylation of its activation loop serines. Additionally, this class of inhibitor partially blocks IKK2’s catalytic activity by enhancing dynamics within the ATP binding pocket and likely the general active site. Hydrogen deuterium exchange (HDX) experiments further revealed that while binding of substrate ATP perturbs only the local structure surrounding its binding site, binding to ATP-competitive or allosteric inhibitors induces structural perturbations in an expansive area including the helical domain. We propose that these allosteric sites can act as specific targets for the development of novel potent IKK inhibitors.

**SIGNIFICANCE:** Aberrant activation of IKK2 is the leading cause of most inflammatory diseases and many associated cancers. Most inhibitors developed for IKK2 are ATP-competitive, and they are toxic *in vivo* due to their off-target effects. By combination of virtual screening and cell-based functional assays, we identified small-molecule binders of the class of benzoyl conjugates of pyrrolidinedione that block IKK2 activity from an allosteric site through covalent attachment and could be specific only for IKK2. HDX-MS and MS-MS assays identified a binding pocket with a ‘Cys-Cys motif’ for these inhibitors, and revealed specific differences in IKK2 dynamics upon binding to substrate ATP vs ATP-competitive and allosteric inhibitors. Present work provides a framework for the development of allosteric inhibitors to combat IKK2-induced diseases inhibitors.

## INTRODUCTION

The NF-κB family of dimeric transcription factors orchestrates elaborate gene expression programs to meet the need for a vast number of cellular functions including innate and adaptive immunity and inflammation (1, 2). In most unstimulated cells NF-κB is inactive. A plethora of stimuli, pathogenic or physiological, acts on specific but diverse receptors, and the resultant signals ultimately converge onto and activate a protein kinase complex known as the inhibitor of κB kinase (IKK). Active IKK phosphorylates IκBα; (inhibitor of κB) of the IκB:NF-κB complexes within which NF-κB dimers are held inactive. Phosphorylation by IKK leads to lysine-48 linked polyubiquitination of IκB followed by its proteasome-mediated degradation and release of free NF-κB dimers for transcriptional regulation (3). In almost all inflammatory diseases including inflammation-induced cancers and neurological disorders, NF-κB is constitutively active (4); and constitutive activation of IKK underlies constitutive NF-κB activity. Mutations in genes encoding factors involved in IKK activation pathways are responsible for pathogenicity, although the vast majority of disease-causing events are sporadic.

The IKK complex is composed of two catalytic subunits, IKK1 (also known as IKKα;) and IKK2 (IKKβ) and an adapter subunit, NEMO (<u>N</u>F-κB <u>E</u>ssential <u>MO</u>dulator) (5, 6). Most NF-κB signaling pathways propagate through the NEMO-IKK2 arm of the heterotrimeric IKK complex activating primarily the NF-κB p50:RelA heterodimer, referred to as canonical signaling. The precise role of IKK1 in this canonical signaling is still debatable. IKK1, however, is an essential mediator of non-canonical signaling which predominantly induces the activation of the NF-κB p52:RelB heterodimers through the processing of the precursor protein p100 protein into p52 (7). The X-ray structures of the core domains of both human IKK1 and IKK2 homodimers are available (8-11). These structures reveal a conserved structural organization where the monomer is folded into four interconnected domains. From N to C, the first three domains are the kinase domain (KD), ubiquitin-like domain (ULD), and a long three helical bundle referred to as the scaffold dimerization domain (SDD). The very last flexible segment of about 85 residues interacts with NEMO, and is known as the NEMO binding domain (NBD). *In vivo*, the IKK complex is predominantly heterotrimeric, i.e. the NBD of both IKK1 and IKK2 subunits contacts the N-terminal domain of NEMO. Even though all three subunits form stable independent homodimers in isolation, the IKK2:NEMO complex is a distributive oligomer. Most likely, the IKK1:IKK2:NEMO complex also forms similar high-order oligomers. It is not known, however, if an oligomeric organization is required for IKK activation. IKK activation is synonymous with the phosphorylation of two serines located within the kinase signature activation loop (AL). Alteration of these serines to phosphomimetic glutamates renders IKK constitutively active *in vivo*. Many kinases, including IKK2 itself, have been shown to phosphorylate the AL of IKK2 during canonical signaling.

For the past 20 years, there has been an intense search for IKK inhibitors for therapeutic application (12). Although many inhibitors have been developed and some of them bind tightly *in vitro*, all failed in clinical trials (13). Most of these small molecule inhibitors are ATP competitive. Toxicity of one of these ATP-competitive inhibitors, MLN120B, has been extensively studied and it has been found to be toxic in animal models (14-17). It is surprising that the development of effective IKK inhibitors is faced with such difficulties, especially when protein kinases have been the most intensely pursued drug targets with >30 FDA-approved drugs and >130 additional compounds in various stages of clinical trials targeting various kinases (18). It is possible that due to the high similarity of the kinase domain with many other kinases (often more than 30 % identity), it is difficult to exploit unique chemical features of the nucleotide-binding pockets of IKK catalytic subunits. Structural analyses of IKK2 reveal the position of an ATP-competitive inhibitor K252a overlapping the binding site of substrate ATP, however the specificity determinants unique to the IKK catalytic pocket have not been explored in-depth and exploited (8). Given the challenges to identifying specific inhibitors for the IKK2 catalytic pocket so far, we employed an alternate approach towards targeting a pocket outside of the active site and obtaining highly specific inhibitors. These types of inhibitors are referred to as class IV inhibitors (19, 20). We particularly targeted development of type IV inhibitors that would block IKK2 activation signaling by modulating interactions between the KD and ULD or SDD. We initially screened for pockets with druggable properties outside the KD of IKK2 that could accommodate small molecules. Depending on the mechanistic properties of the IKK complex architecture, these binders may or may not block IKK2 kinase activity to phosphorylate substrates *in vitro*. We adopted virtual docking for preliminary identification of binders to these pockets separated from the ATP-binding pocket. We tested the top 100 commercially available compounds and confirmed two related compounds as inhibitory hits for IKK2 activation using cell-based experiments. Further studies using HDX-MS and MS-MS assays enabled identification of the binding pocket for these compounds outside of the ATP binding site. The binding site pocket is located at the SDD-KD junction where a Cys-Cys motif within the kinase domain is covalently modified by the electrophilic inhibitors. HDX-MS studies reveal that the inhibitor binding induces flexibility of the ATP binding pocket suggesting a communication between the ATP binding pocket and the inhibitor binding site. Furthermore, both ATP-competitive and allosteric inhibitors induce similar protection or flexibility outside of their binding site in contrast to ATP that affects only its binding site in the kinase domain. Overall, these experiments result in the development of high-affinity ligands targeting an allosteric site separated from the catalytic pocket and provide us with a conceptual platform to discover novel therapeutic agents.

## RESULTS

### The nucleotide binding pocket of IKK2

We wished to investigate why screened IKK inhibitors are particularly targeted to the nucleotide-binding site, and if the lack of uniqueness of the ATP-binding pocket posed challenges in obtaining IKK-specific ATP-competitive inhibitors. There is one reported structure of human IKK2 bound to staurosporine analog K252a providing information on how ATP-competitive inhibitors could recognize IKK2 (pdb: 4KIK)(8). Efforts from our laboratory provided a modest resolution crystallographic model of human IKK2EE (two serine residues of the AL mutated to glutamate) (pdb: 4E3C) (9). This provided several key structural insights, however, even though these crystals were grown in the presence of an inhibitor (MLN120B), the reported structural coordinates did not locate the inhibitor. We revisited the refinement to see whether we could improve upon the maps to observe density of MLN120B in any of the six IKK2 molecules (three reciprocally stacked dimers) in the asymmetric unit. The deposited structure factor and phase data of pdb id 4E3C was used to further refine the map and coordinates using programs REFMAC5 and Coot, respectively. Indeed, we found densities accounting for MLN120b bound to the nucleotide-binding pocket in three of six protomers (**Figure 1A**).

**Figure 1.**
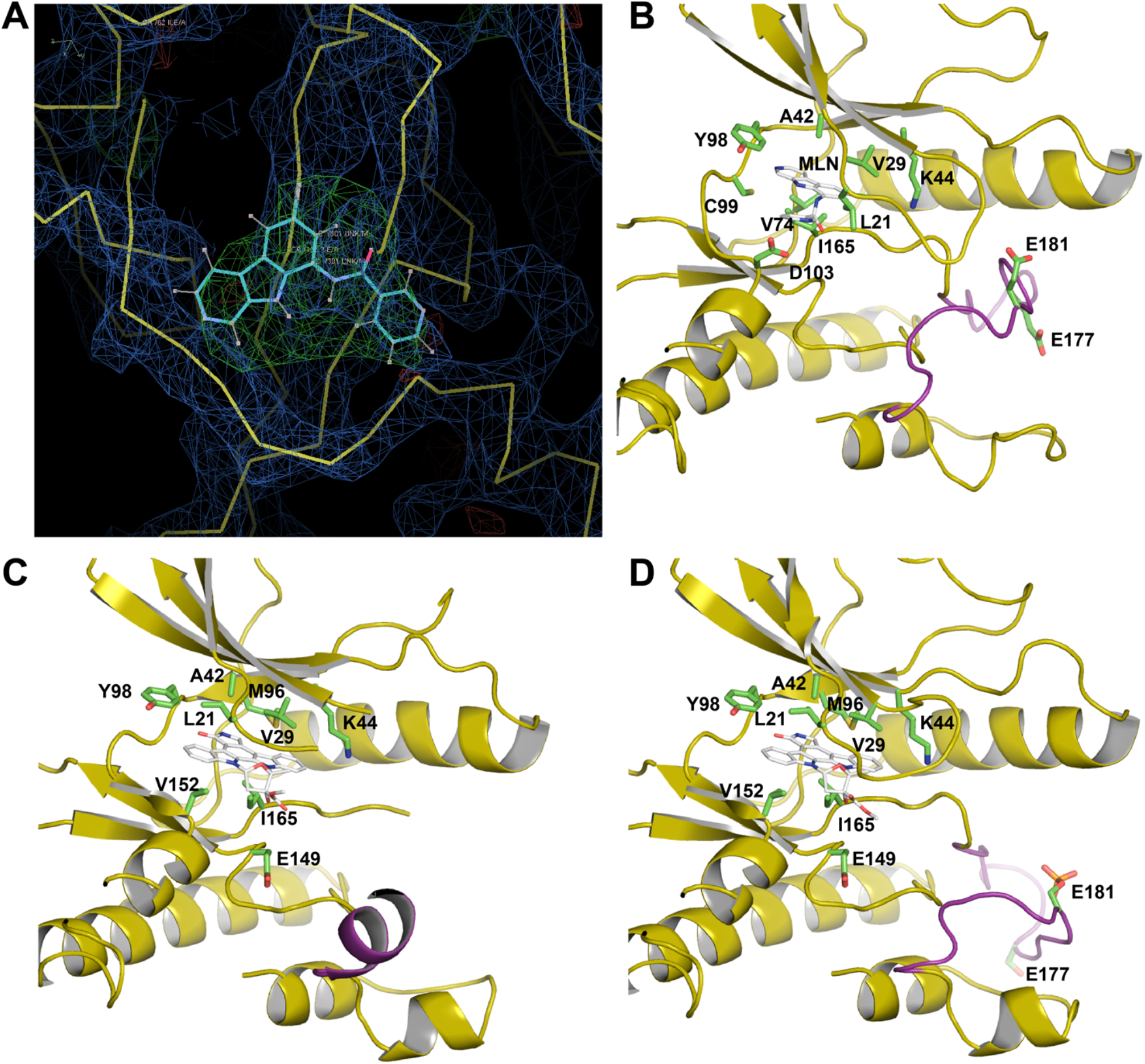
Binding of inhibitors to IKK2 active site. A. A difference density map at sigma contour level of 3 showing the position of inhibitor MLN120B in the active site pocket of IKK2. B. A cartoon diagram showing interaction of primarily hydrophobic residues with MLN120B in the pocket. C. Inactive IKK2 protomer with flexible AL from 4KIK bound to K252a. D. Active IKK2 protomer with structured AL from 4KIK bound to K252a, AL is structured differently from that of IKK2 monomer bound to MLN120B.

The occupancy of the inhibitor varies, nonetheless the general area of the inhibitor is clear, especially in one IKK2 monomer. A comparison of the binding pocket of MLN120B to that of staurosporine analog K252a in human IKK2 (8) reveals a general overlap of the binding location, and naturally many common residues e.g. Leu21, Val29, Ile165 are engaging in the interactions (**Figure 1B**). Nonetheless, the binding is specific with MLN120b engaging different residues such as V74 and D103. The binding mode of MLN120B to independent hIKK2 protomers appears similar. In the lattice of the 4E3C crystal, all six protomers appear to be in the active conformation, although ALs display variable flexibility and this flexibility shows a reverse correlation to the occupancy of the MLN120B, perhaps as anticipated. Conversely, in the crystal lattice of the 4KIK structure, one subunit is in the active conformation and the other is in the inactive conformation although both subunits of the dimer engage to the inhibitor nearly identically (**Figure 1C, D**). Notably, the AL conformation of the protomer in the active state in 4KIK differs from that of the AL of active state monomers in 4E3C (**Figure 1B, D**). Since the class I inhibitors MLN120B and K252a bind similarly to subunits of different conformational states, these do not appear to discriminate between the structural states i.e., functional activation states of the kinase domain. It is unclear why class I inhibitors of IKK were preferentially identified during screening and why these class I inhibitors were toxic and therefore not useful as therapeutics (13). Since massive chemical screening efforts have been conducted and over 130 patented IKK inhibitors exist, it could be argued that typical screening strategies are perhaps not ideal for identification of IKK inhibitors appropriate for therapy. Screening with alternate approaches to identify other classes of IKK kinase inhibitors for drug development has not been performed with rigor.

### Identification of allosteric sites by virtual screening

Mutagenesis experiments indicated several protein-protein interfaces in addition to the primary IKK2 dimerization surface (SDD-SDD) to be important for IKK2 activation (9). One of these trans-dimer interfaces, named the ‘V’ interaction surface, contained residues I413, L414, P417 and K418, and mutations of these residues to alanine reduced IKK2 activation ex-vivo suggesting a possible role of this V surface. Based on this information, we searched for small molecule binders to the V surface area, and other pockets away from the nucleotide-binding pocket that can prevent canonical signaling. We used virtual screening (VS) around the center of mass of residues I413, L414, P417 and K418 in pdb ID:4E3C. Using Schrodinger’s Glide program (21) we docked into this site a select in-house library of approximately 1800 compounds in ChemBridge database accessible to us, and ranked binding by the ligand efficiencies, which is equivalent to the binding score divided by the heavy atom count (described in the methods section). Binding scores were calculated using Glide’s empirical scoring function that approximates the ligand binding free energy. In addition to the top-scoring 10 compounds in ligand efficiency, 90 additional compounds above the median of the ligand efficiency were also selected to be tested for IKK2 inhibition (**Supplementary Table**).

### Ligands blocking IKK2 activation in cells

We performed cell-based assays to determine if any of the 100 compounds selected based on virtual screening could inhibit NF-κB activation in model mouse embryonic fibroblasts (MEF) cells (Figure 2A-D). We monitored degradation of IκBα and phosphorylation of IKK activation loop by Western blot (WB) assay with appropriate antibodies. In addition, we monitored phosphorylation of IκBα by WB and specific DNA-binding efficiency of RelA dimers of the NF-κB family using electrophoretic mobility shift assay (EMSA). One compound among the 10 top-scoring compounds (cpd 4), and one among the rest 90-compound set (cpd 65), were found to prevent degradation of IκBα signifying blockage of IKK2 activation pathway intermediates (**Figure 2A, 2B**).

**Figure 2.**
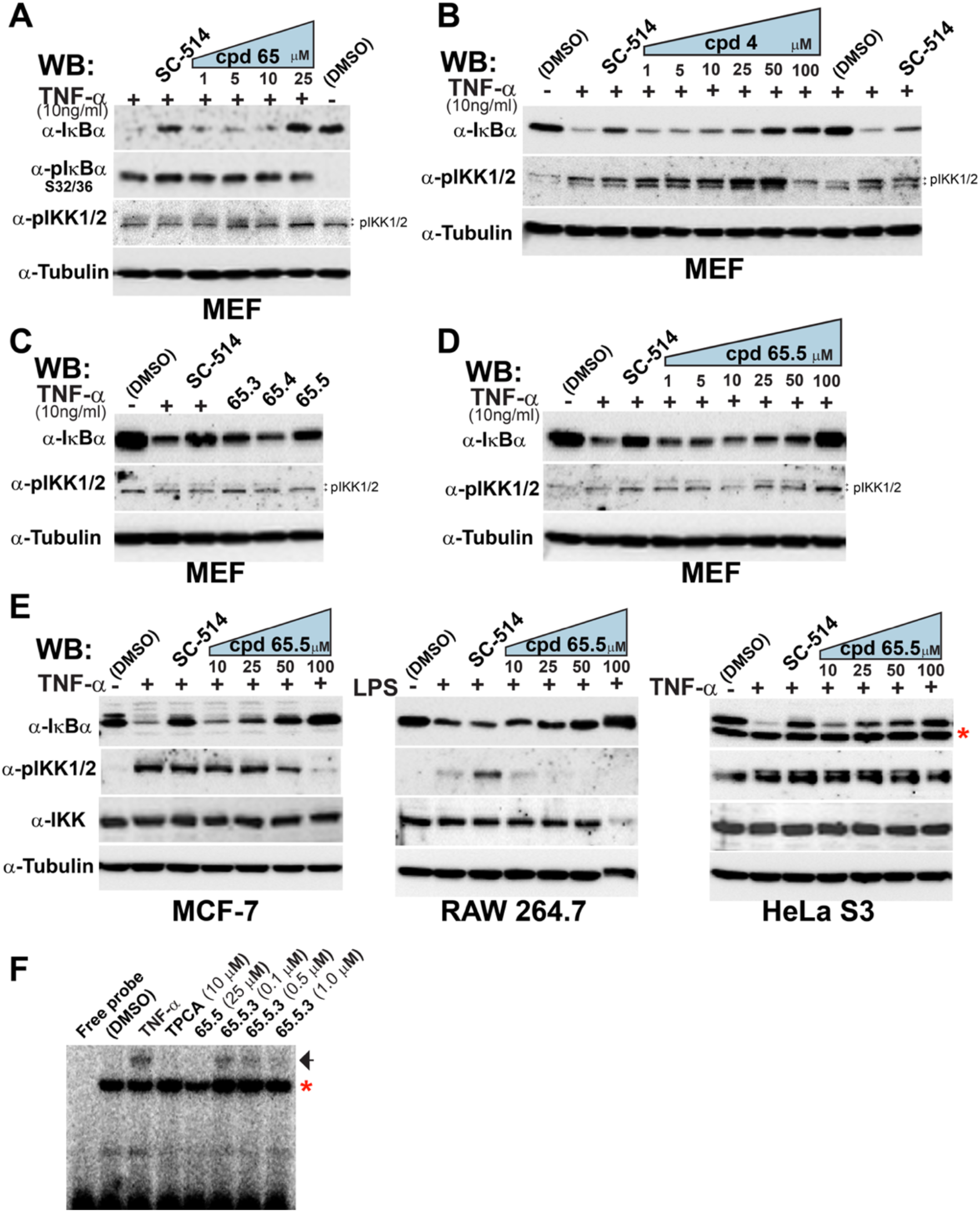
Effect of inhibitors on IκBα, phospho-IκBα, phospho-IKK, and NF-κB DNA-binding level in TNF-α treated cells. pIKK antibody detects two bands: pIKK2 (upper), and pIKK1(lower). **A**. MEF cells were stimulated with TNF-α in presence of IKK inhibitor SC-514 or increasing concentrations of cpd 65. **B**. MEF cells were treated as indicated in section A in presence of increasing concentrations of cpd 4. **C**. MEF cells were treated as indicated in section A in presence of 10 μM of compounds 65.3, 65.4 and 65.5. **D**. MEF cells treated as indicated in section A in presence of increasing concentrations of cpd 65.5. **E**. MCF-7, RAW 264.7 and HeLa S3 cells were stimulated with TNF-α; or LPS in presence of SC-514 or increasing concentrations of cpd 65.5. Red star indicates the presence of a non-specific band in HeLa S3 extract. **F**. EMSA showing level of endogenous nuclear RelA by its binding to HIV-κB DNA in TNF-α; stimulated MCF7 cells in presence of IKK2 inhibitor TPCA or cpds 65.5 and 65.5.3. Single black arrow indicates the specific NF-κB-DNA complex. Red star indicates a non-specific band.

TPCA(22) and SC514(15), the ATP analogs used as positive controls of IKK2 inhibition, blocked IκBα degradation but failed to block IKK2 activation as reflected in absence of pIKK reduction. In contrast, cpd 65 blocked both IκBα degradation and phosphorylation of the activation loop of IKK2 activation (upper band of the phospho-IKK doublet in figures 2A-2D). The inhibition of IKK AL phosphorylation correlated well with IκBα degradation. These observations hints cpd 65 to be non-ATP competitive and blocking IKK2 activation allosterically thereby a kinase inhibitor of class IV.

In contrast to cpd 65, cpd 4 promotes the activation loop phosphorylation of both IKK1 (corresponds to the lower band of the doublet) and IKK2 indicating its broader binding specificity to both IKK subunits (**Figure 2B**). It is possible that cpd 4 binds to a site that is common to both subunits. Since our focus is to identify IKK2-specific allosteric inhibitors, we expanded the structure-activity relationship (SAR) analysis via second-generation derivatives of cpd 65. We tested 13 commercially available compounds related to cpd 65 (cpds 65.1 to 65.13) (**Supplementary Figure 1A**), and identified two compounds (cpd 65.3 and cpd 65.5) (**Supplementary Figure 1B**) with an IC50 of ∼10 μM i.e., slightly improved inhibitory potency compared to cpd 65 (**Figure 2C, Supplementary Figure 1C**). Since both these compounds had similar properties, follow-up experiments were performed with only cpd 65.5 in HeLa, RAW, and MCF7 cell lines (**Figure 2E**). IκBα degradation was inhibited in all cells although strength of inhibition evaluated from level of IκBα degradation (and IKK2 AL phosphorylation) is different in different cell lines. Significantly higher concentrations of cpd 65.5 are required for inhibition in the cancer cell lines than in MEF. At higher concentrations (> 50 μM), cpd 65.5 appeared to be toxic to RAW cells. We further tested NF-κB activation by assessing DNA binding level of endogenous NF-κB RelA dimers in MEF cells stimulated with TNF-α in the presence and absence of cpd 65.5 using EMSA. As anticipated, cpd 65.5 reduced level of specific DNA bound to NF-κB RelA dimers in MEF cells stimulated with TNF-α (**Figure 2F**).

Since cpd 65 and cpd 65.5 contain highly electrophilic carbons, we reasoned that these inhibitors might be functioning through covalent linkage to cysteines. To test this, we generated a more electrophilic compound (cpd 65.5.3) and tested its inhibitory properties in MEF. The results suggest that specific binding and covalent modification properties are both essential in blocking IKK2.

We also tested if the inhibitors block non-canonical NF-κB signaling which is distinct from canonical signaling to assess their specificity. Ligands signaling Non-canonical pathway, distinct from canonical signaling pathway, trigger NF-κB precursor protein p100 to be processed to p52. IKK2 has no reported role in non-canonical signaling. Cpd 65.5 treatment had no discernible effect on p100 processing in MEF cells suggesting its inhibitory specificity for only canonical signaling (**Supplementary Figure 1D**).

### Dynamic segments of IKK2 11-669 EE map into the loop and linker regions

We used hydrogen-deuterium exchange coupled to mass-spectrometry (HDX-MS) experiments (23) to determine the putative binding sites of inhibitors of IKK2. First, we determined the deuterium uptake into free IKK2. For these experiments, we used IKK2 11-669 EE since this form was used for crystallographic structure determination. 135 peptides that covered 85.1 % of the IKK2 (11-669 EE) sequence could be separated using their size, mobility, and retention time on a liquid chromatography column and identified/analyzed using Waters MassLynx software.

Areas of high exchange of hydrogen with deuterium indicate regions of the protein that are solvent-exposed. Areas of low exchange indicate more tightly folded regions of the protein with less exposure to solvent (**Figure 3A**).

**Figure 3.**
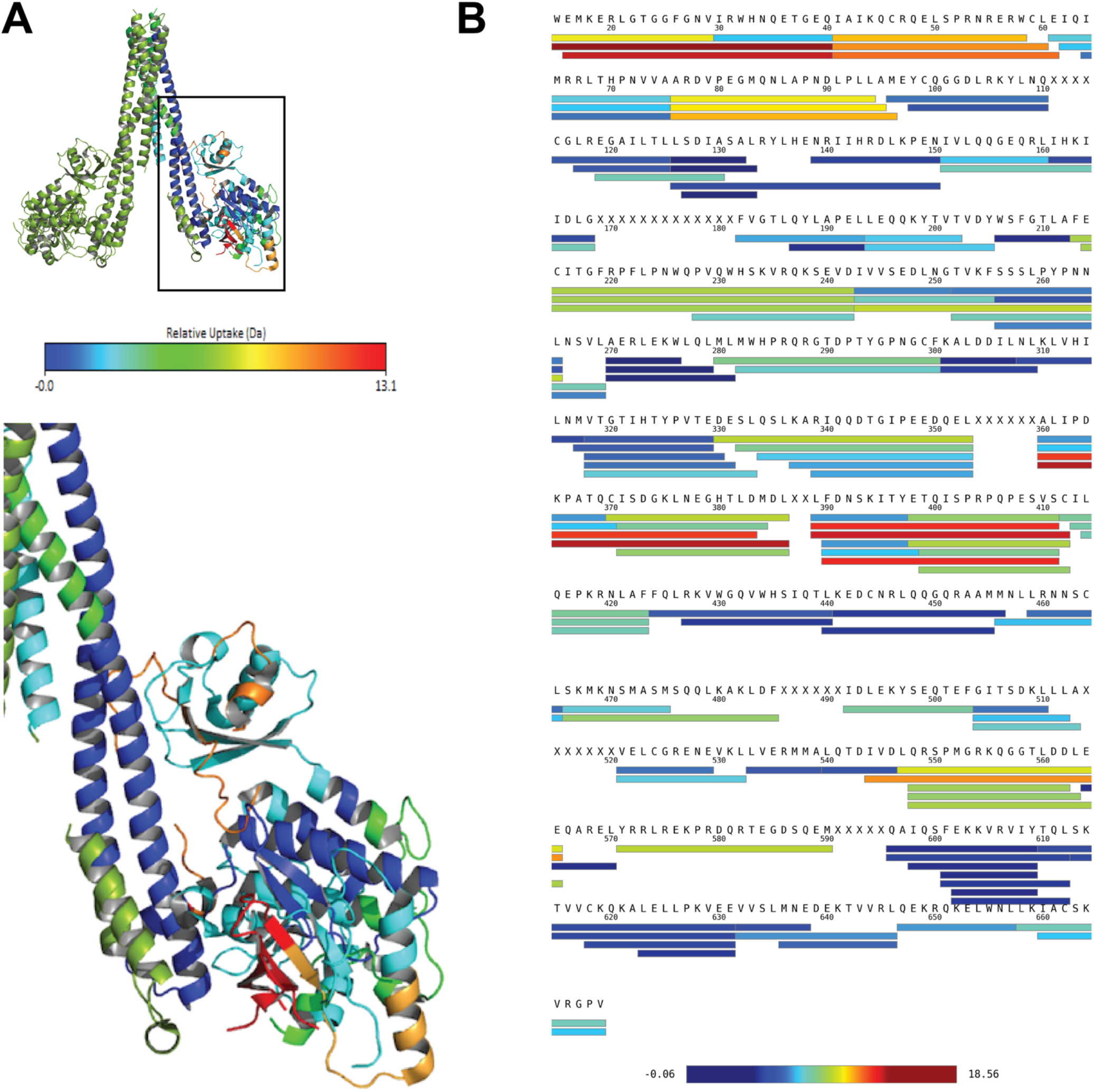
Dynamics of IKK2 assessed by HDX-MS experiments. Rainbow spectrum indicates the spread of exchange with dark blue indicating areas of minimal exchange, and red areas of highest exchange (A) Exchange mapped IKK2 dimeric structure, only a monomer is highlighted for exchange and an inset shows the magnified view of the junction of KD, ULD, and SDD. (B) Exchange mapped on the primary sequence.

The HDX data indicates areas of high flexibility within the kinase domain, particularly residues encompassed within residue range 15-40, and within residue ranges 41-61 and 75-95 (**Figure 3B**). Residues contained within ranges 125-150, 205-210, 270-280, and 300-310 of the kinase domain exchanged less, presumably because these are well-folded/less dynamic. We also observed high flexibility in another area spanning residues 360 to 410 i.e. located primarily in the ubiquitin-like domain (ULD) but crossing over into the scaffold dimerization domain (SDD) (**Figure 3A**). Overall, the ULD indicated an intermediate level of exchange throughout with no region exchanging as little as the well-structured parts of the kinase domain (**Figure 3A, B**). The SDD exchanged the least of the three domains with little exchange in residues within 427-455, 563-570, and 595-657 (**Figure 3A, B**). Interestingly, residues within SDD regions 545-563, and 570-590 exchanged highly even though these are next to the low exchanging region 563-570 (**Figure 3A, B**).

### IKK2 behaves differently upon binding to ATP versus inhibitors

We next tested HDX of IKK2 in the presence of inhibitory cpd 65.5.3. These experiments were performed in two sets; in first set, we compared HDX of IKK2 in free, ATP-bound, ATP-competitive inhibitor TPCA-bound (22, 24), and cpd 65.5.3-bound states. In second set, we compared HDX of IKK2 in free and cpd 65.5-bound states (**Figures 4A-G**).

**Figure 4.**
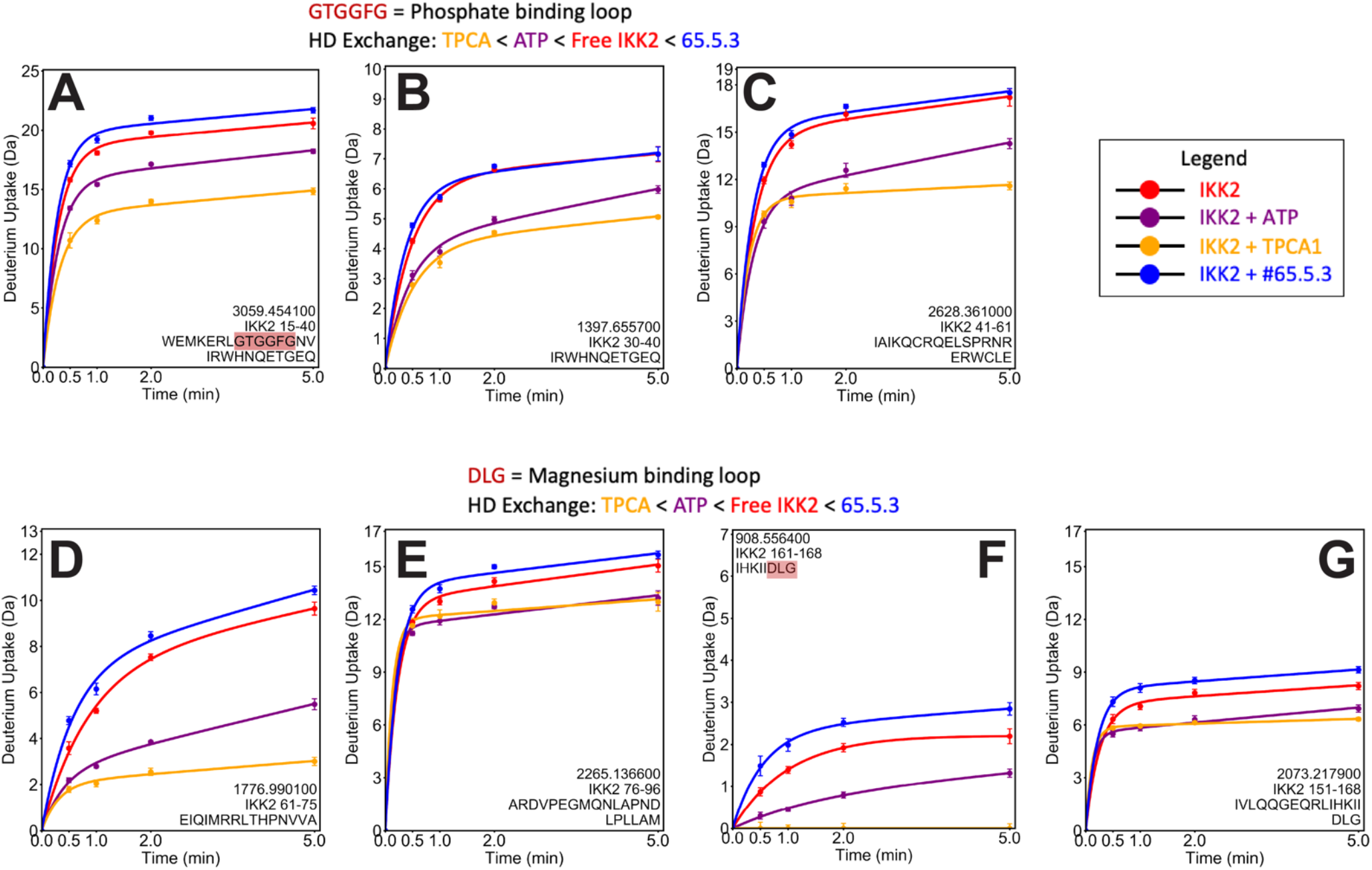
HDX-MS profiles showing comparison of the protection and deprotection of H-D exchange in IKK2 in its free (red), partnered with ATP (purple), ATP-competitive inhibitor TPCA (orange), and cpd 65.5.3 (blue) states. Treatment with cpd 65.5.3 displays deprotection in the nucleotide phosphate-binding loop (**A, B, C**) and in the Mg binding loop (**D, E, F, G**) and surrounding peptides while treatment with ATP and TPCA show protection.

A decrease or increase of exchange in areas observed upon binding of ligands when compared with that of free protein might be indicative of possible areas of ligand interaction. Decrease or increase in exchange could also be reflective of alteration in dynamics due to allosteric effects as a result of ligand binding. We refer to a decrease in exchange as the ‘protecting’ behavior. In contrast, an increase in exchange could occur as a result of increased dynamicity and/or solvent exposure upon binding the partner, and referred to as the ‘deprotecting’ behavior.

We first analyzed the kinase-conserved nucleotide-binding segments 15-100 and 150-170, which include the Mg-binding loop. Conserved residues that are critical for binding to ATP are all contained within these regions. These regions include the glycine loop ^22^GTGGFG^27^ that contact the phosphate of ATP, K44 that contacts the α and β phosphate of ATP, E61 that stabilizes K44 by forming a salt bridge, hinge residues E97 and C99 that bind to N6 of adenine, and residues L21, V29, A42, M96, Y98, and V152 that mediate hydrophobic contacts. The most striking HDX differences are observed in residues of pocket encompassing stretches 15-40, 41-61, 61-75, 76-96 and 151-168 (**Figures 4A-G**). No ATP-bound structure of IKK2 is available, nonetheless protection data of HDX-MS clearly suggests that IKK2 uses all predictive signature sequence features in binding ATP. Protection upon binding to ATP and TPCA in these peptides and its shorter segments suggests that both engage a similar binding pocket. However, TPCA induced a stronger protection compared to ATP, consistent with its much stronger binding affinity. This is also supported by the observation that protection of peptides 15-29 and 96-110, both regions located within the ATP binding pocket, is clearly observed upon binding to only TPCA but not ATP (**Supplementary Figure 2A, Panels 1, 14**). A surprising distinction in ATP vs TPCA mediated protection is that ATP does not show any deprotection within the kinase domain or in the rest of the kinase molecule, whereas, several regions of the full kinase and two regions, residues 213-242 and 243-265, within the kinase domain in particular, are deprotected by TPCA. In contrast to ATP and TPCA, cpd 65.5.3 did not show protection of any of the peptide segments in the nucleotide binding pocket (**Figure 4)**. This is consistent with our cell-based experimental data showed cpd 65.5.3 bound a region outside the nucleotide binding pocket, and was not an ATP-competitive inhibitor. Notably, the enhancement of deprotection at the ATP-binding pocket compared to unliganded kinase suggests that the binding of cpd 65.5.3 modulates IKK2 allosterically. These results classify cpd 65.5.3 as a class III or IV kinase inhibitor. Similar to TPCA, cpd 65.5.3 also deprotects residues of two other regions, 213-242 and 243-265, which partly overlaps helices αF and αG and their connecting segment within the kinase domain similar to TPCA (**Supplementary Figure 2B, Panels 12, 18**). The first peptide interacts with SDD and the latter with ULD.

### Both TPCA and cpd 65.5.3 deprotect several common regions outside of the kinase domain

The ULD region did not display any significant difference in deuterium uptake upon binding to inhibitors or ATP with the exception of peptide spanning residues 360-386 and surrounding peptides. These peptides show deprotection upon cpd 65.5.3 treatment (**Supplementary Figure 2D, Panel 3, 4, 6, 7**). Surprisingly, cpd 65.5.3 and TPCA show a remarkably similar deprotection pattern in several areas within the SDD that include four peptides spanning residues 466-485, 548-563, 571-590, and 632-646. ATP showed no discernible effect in these regions (**Supplementary Figure 2E, Panels 8, 20 and Supplementary Figure 2F, Panels 3, 17**). When the deprotection from the kinase domain is considered, TPCA and cpd 65.5.3 both bring similar allosteric structural change at regions distal to their binding sites spread over seven peptides. The significance of this similarity in molecular dynamics experienced by the kinase upon binding to two different types of inhibitors occupying distinct sites is unclear. Nonetheless, binding of TPCA appears to propagate dynamicity resulting in both a strong protection in areas close to the binding site and deprotection at sites away from the binding site. ATP binding results primarily in protection of the binding site region with no discernible effect outside the KD, suggesting that binding of ATP is coordinated locally with minor if any effects of distal regions. By extension, it can also be suggested that during substrate phosphorylation, ATP binding and activity has evolved to be restricted to the kinase domain. Synthetic inhibitors are not under any similar constraints, and thus upon binding could trigger disconcerted molecular motions.

### Cpd 65.5.3 and cpd 65.5 bind to a common site

Cpd 65.5.3 showed no protection in any of the domains of IKK2 with the exception of segment spanning residues 115-125 (**Figures 5A, B**).

**Figure 5.**
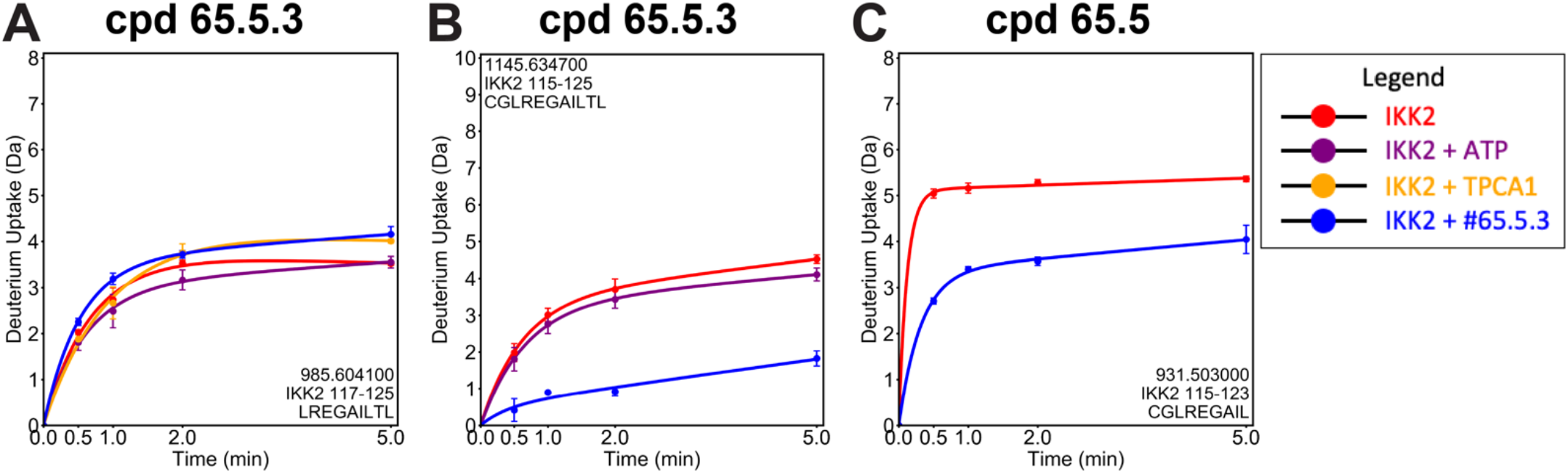
IKK2 peptides protected by inhibitors 65.5.3 and 65.5 from H-D exchange. Compounds 65.5.3 and 65.5 show deprotection in overlapping peptides 115 to 125 (**B**) and 115 to 123 (**C**), respectively. These peptides include the cysteine 115 of the di-Cys 114/115 motif. Comparison of behavior of fragment 117-125 without cysteine (**A**) to 115-125 (**B**) with compound 65.5.3 indicates cysteine to play important role in binding of 65.5.3.

Cpd 65.5 also showed protection of this region (**Figure 5C**). Interestingly, peptide spanning residues 117-125 showed no protection when bound to 65.5.3. This suggests interaction between the kinase and 65.5.3 occuring through one or both of the two additional residues. One of those residues is cysteine 115 and the other is glycine 116. Cysteine 115 is part of a di-cys motif with an additional cysteine located at position 114. We speculated that cpds 65.5 and 65.5.3 might be covalently linked to these cysteines. This segment is insensitive to exchange upon binding of both ATP and TPCA. These two cysteines contact the SDD and a close examination of these cysteines in structural models reveal that they could form a disulfide bond. These two cysteines were also identified to be extremely sensitive to oxidation, and IKK2 is inactivated upon their oxidation (25). However, a lack of a protection of segments nearby Cys114/115 suggests that binding of cpds 65.5 and 65.5.3 are unlikely to enforce any elaborate non-covalent contacts with other residues and their primary mode of inhibition is through the oxidation of Cys114/115.

### Both cpd 5.5 and cpd 65.5.3 form covalent bonds with Cys114 and Cys115

To test if cpds 65.5 and 65.5.3 bind covalently to IKK2 11-669 EE, we incubated them individually with the kinase and subjected the reaction mixtures for MS-MS analysis. MS-MS data indicates peptides with altered masses in cpd 65.5 and 65.5.3-treated proteins reflecting likely mass increases on cysteines 114 and/or 115 (**Figure 6**).

**Figure 6.**
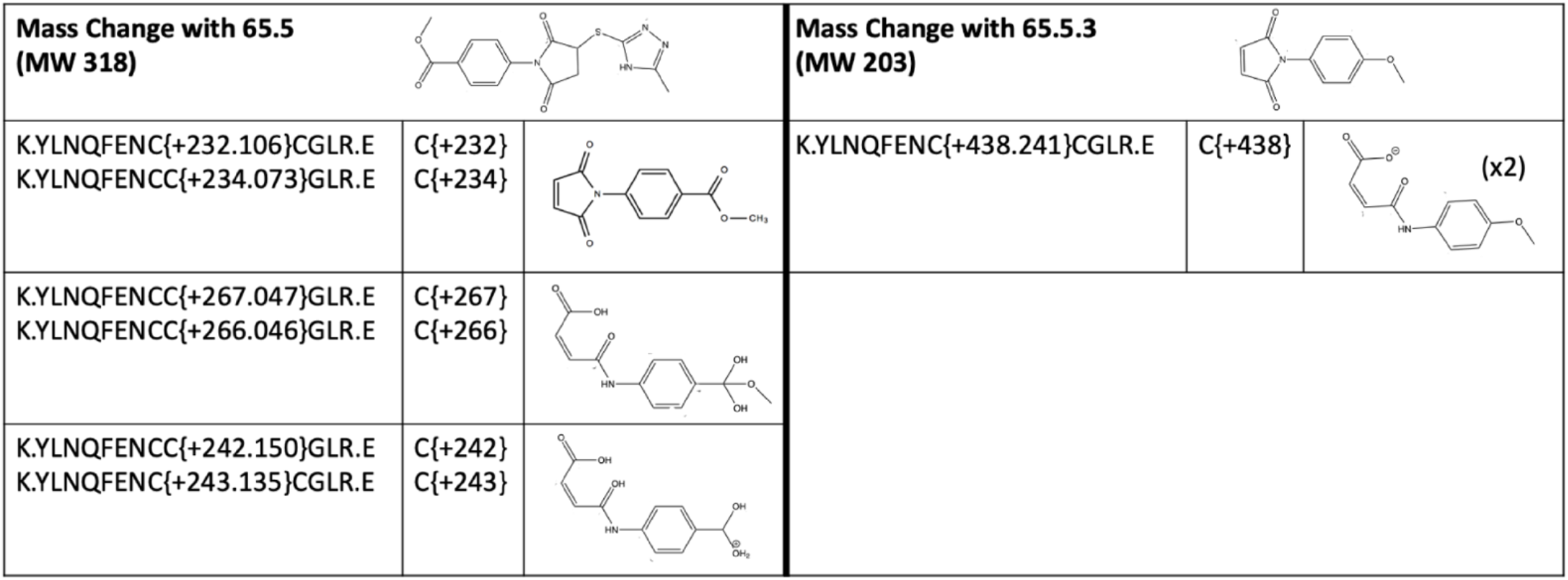
MS-MS data showing inhibitors bound to Cys114 and Cys115. IKK2 was incubated with inhibitors for 30 minutes on ice before separation on a 10 % SDS PAGE gel. The MS-MS analysis of IKK2 bands excised from the gel indicates alteration of mass of peptide fragments containing cysteines 114 and/or 115. Altered masses are indicative of reaction with elimination and hydrolysis reaction products of cpds 65.5 and 65.5.3.

Detailed examination indicated that different masses are a result of multiple reaction products of the inhibitor undergoing β-elimination and hydrolysis reactions. We observed only a single peptide adduct from the reaction with 65.5.3, and the MW of the adduct suggested hydrolytic products of two 65.5.3 molecules reacted with both the cysteines 114 and 115 of IKK2. Comparison of data from HDX-MS runs with and without DTT also confirmed the inhibitor binding activity in segment 115-125 (**Figure 5B, Supplementary Figure 2A panel 16, Supplementary Figure 3A**). No protection of this segment was observed when the inhibitor was added in the absence of DTT. This suggests that Cys114 and Cys115 are likely primarily in their oxidized state preventing inhibitors from forming a covalent bond. In the crystal lattice of inhibitor bound 4KIK Cys114/Cys115 of IKK2 are not disulfide bonded reflecting that in the inhibitor bound form both Cys114 and Cys115 of IKK stay reduced (**Supplementary Figure 3B, C**). Overall, both HDX and MS-MS data reveal that the electrophilic inhibitors covalently couple to this highly oxidation sensitive di-Cys motif and trigger effects on residues of distant areas including ATP-binding pocket and thus cause inactivity of IKK2.

### ATP-binding and substrate phosphorylation activities of active IKK are partially inhibited by cpds 65.5 and 65.5.3

Our experiments indicated cpds 65.5 and 65.5.3 bind to a site distant from the ATP-binding pocket and prevent activation of IKK2. However, these inhibitors might not inactivate already activated IKK2. We performed an in vitro kinase activity assay with Sf9-derived IKK2 11-669 EE that is already active to test if cpd 65.5 could inhibit its kinase activity. As a control, we used SC514, TPCA, and MLN120b, all ATP competitive inhibitors. As anticipated, these ATP-competitive inhibitors inhibited substrate IκBα phosphorylation, but cpd 65.5 did so only partially (**Figure 7A** and **Supplementary Figure 4A, B**).

**Figure 7.**
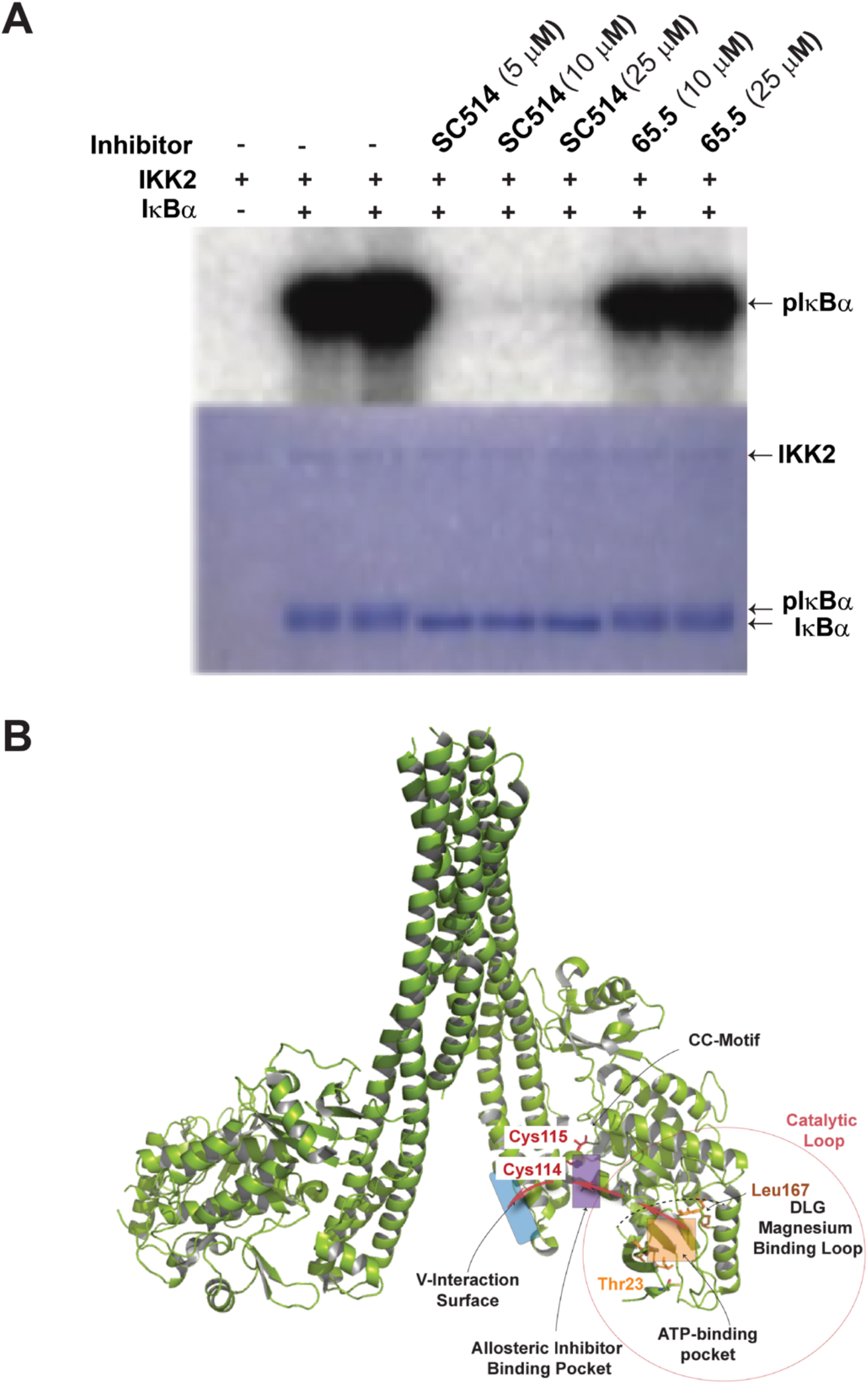
Inhibitory effects of SC514 and cpd 65.5 assessed by phosphorylation of IκBα. **A**. Reaction mixtures (100 ng IKK2, 200 μM ATP, 10 μCi ^32^P-labeled ATP, 1 μg IκBα) were incubated with inhibitors for 30 minutes at RT, quenched with SDS PAGE dye, separated in SDS-PAGE, stained with Coomassie (bottom), and autoradiographed. Phosphorylation of IκBα by IKK2 is nearly abolished by SC514 and significantly reduced with cpd 65.5. **B**. A schematic of IKK2 with the ATP-binding pocket (orange), inhibitor binding pocket (purple) with the two cysteines (red), Mg-binding loop, and the V-interaction surface (blue) highlighted. Thick red arrows indicate the imaginary path of allostery.

This partial inhibition of activity of cpd 65.5 is consistent with our HDX-MS data that indicated enhanced flexibility of the ATP binding pocket, which likely correlates with inhibition (partial) of ATP binding. We observed similar results with cpd 65.5.3. In contrast, MLN120b nearly abolished IKK2 catalytic activity. These results strongly suggest that cpds 65.5 and 65.5.3 function as allosteric inhibitors and not as ATP-competitive inhibitors of IKK2 activation.

## DISCUSSION

The focus of our study is to identify specific pockets of IKK2 besides its ATP-binding pocket that contain residues critical to its activation, and thus could be targeted by small-molecule inhibitors. A couple of recent reports described two inhibitors of IKK2 that might also function allosterically. One of these inhibitors is postulated to interact with IKK2 at the interface between ULD and KD based on modeling experiments but its proof awaits experimental evidence (26). The other inhibitor, a natural product, binds within a small lobe of the kinase domain and covalently couples to Cys46 (27). Perhaps by docking to the small lobe, the inhibitor affects the dynamic structure of the KD and results in allosteric inhibition of IKK2 activity. Efficacy of these allosteric inhibitors as drugs requires further investigation. We surmised the feasibility of finding many such pockets since IKK2 is a large protein of several domains, and a well-knit juxtaposition of these domains is necessary for both its activation and for the catalytic activity of the kinase domain. We previously observed that a monomeric IKK devoid of its accessory domain ULD and a large part of the SDD is incompetent in phosphorylating its substrate IκBα at two N-terminal serines that is required for IκBα degradation - even if the two AL serine residues of this monomeric IKK are changed to phosphomimetic glutamates (28). Moreover, alteration of residues at the domain-domain interface inactivates IKK2. These facts suggest that coordination between independent domains of IKK2 is necessary for its activation and activity (9), and prompted us to look for small molecule binding pockets at the interdomain or dimer-dimer interaction interfaces. Molecules binding to these unique pockets separated from the nucleotide-binding site/catalytic domain will belong to kinase inhibitors of class III and IV and are more likely to possess higher specificity. We first searched for such pockets containing features for small molecule binding, identified a site at the interface of the KD and SDD, and screened for molecules that fit this site using virtual docking. Cell-based experiments confirmed some of these predicted binders to block NF-κB activation in multiple cell lines including cancer cell lines albeit with varied potency. Notably, we found that these inhibitors block IKK2 activation by inhibiting phosphorylation of the activation loop serines in a concentration dependent fashion. In vitro experiments suggested these inhibitors function allosterically since these inhibitors could only minimally block substrate phosphorylation by active IKK2 contrary to a strong inhibition usually observed with ATP-competitive inhibitors. Small molecules targeting this or similar allosteric pockets with high specificity and affinity have the potential to alleviate long-standing challenges faced in identifying specific IKK2 inhibitors (29).

To further explore the mechanism of inhibition, and the site of inhibitor binding, we conducted HDX-MS experiments using IKK2 in free state or bound to ATP, TPCA, and two related allosteric inhibitors identified in this study. We observed that the inhibitory compounds, both containing an electrophilic maleimide moiety, coupled covalently to two cysteine residues (Cys114 and Cys115) and that the inhibition is due to the modification of these two cysteines. Since the benzoyl groups contributed only marginally to the affinity, compounds 65.5 and 65.5.3 on their own are not anticipated to be specific or useful for therapy. Nonetheless, inhibitors targeting this, or a site of similar functionality could meet therapeutic goals. The phenomenon of high reactivity of only Cys114 and Cys115 is somewhat surprising and unexplained, since IKK2 has 9 other cysteines with some others exposed to the solvent. In addition, Cys114 and Cys115 is in close contact with SDD. Cys114 and Cys115 were also reported to be susceptible to mild oxidizing agents such as HOCl, and since such agents are known to be useful in treating skin inflammation (25) it is possible that IKK2 might have a role in this. However, Cys to Ser mutations at 114 and 115 did not affect the catalytic activity; perhaps Ser is not bulky enough to perturb domain-domain interaction and affect IKK activation and activity.

Another puzzling observation is that the location of the bound inhibitors as identified by HDX-MS and MS-MS experiments differed from that predicted by the docking experiment. The CC-motif to which the inhibitory compounds are covalently linked is located next to an open pocket docking site predicted by the docking experiments. The CC-motif and the open pocket are separated by two side chains, thus for cpd moieties to have access intervening residues must shift. Interestingly, the MS-MS experiments indicate that the moieties covalently linked to IKK2 are reaction products of cpds 65.5 and 65.5.3, and not the original compound. Cpd 65.5 underwent a beta elimination of the thiol moiety resulting in a compound with more reactivity toward cysteine. It is possible that screened compounds in their intact forms bind optimally to a specific pocket and induce a structural change employing all their moieties, and consequently generate access to the cysteines for covalent bonding with themselves or their reactive products. Therefore, both the open pocket and the CC-motif might provide optimum substrate engagement capabilities in different steps.

HDX-MS data also revealed some intriguing information related to IKK2 dynamics upon inhibitor binding. ATP induced significant structural changes, although these changes are localized primarily within the kinase domain at the ATP binding site and overall active site. These changes are primarily reduction of the dynamic nature of active site. Somewhat similar effects are observed in other kinases, thus a similar phenomenon could be reflective of a set of ATP-induced structural changes that are conserved and necessary for catalysis. TPCA binds to a pocket overlapping the binding pocket of ATP; however, it induces two opposite changes within the kinase domain, protecting segments surrounding its binding site and simultaneously further exposing (deprotecting) some distal sites to solvent. Furthermore, TPCA influences IKK2 structure beyond its kinase domain, affecting the ULD and SDD. Covalent inhibitors cpds 65.5 and 65.5.3 primarily induced greater flexibility in both the kinase domain and the accessory domains while inducing reduced dynamics at the ATP binding site. The similarity in allosteric structural changes induced by TPCA and the cpd 65 class of allosteric inhibitors suggests an underlying mechanistic phenomenon of communication between the active site and distant allosteric sites (**Figure 7B**). It appears possible to target and disrupt such communication from allosteric sites using small molecules such as the cpd 65 class of inhibitors, and this might allow IKK2 specific inhibition.

## MATERIALS & METHODS

### Western Blot

For IKK2 Inhibition assays, described cells were cultured overnight in 500 μL in DMEM media supplemented with 1 % PSG and 10 % FBS at 37 °C and 5 % CO_2_ in 24-well plates. The cells were incubated first with indicated concentrations of inhibitor or DMSO (as a control) for 2 hours at 37 °C and 5 % CO_2_. TNF-α; was then added at a concentration of 10 ng/mL and the cells were incubated further at 37 °C and 5 % CO_2_ for 15 minutes. Cells were harvested by washing with PBS and lysed in plate with RIPA buffer. The samples were separated on 10 % SDS PAGE gels followed by Western blot analysis using indicated antibodies. The IκBα antibody was purchased from Santa Cruz (Cat # SC-1643), IKK antibody from Millipore (Cat # 07-1479), pIKK antibody from Cell Signaling (Cat # 2694), and Tubulin antibody was a gift from BioBharati Life Science (Kolkata, India).

### EMSA

EMSA assays were performed as previously described(30). Briefly, HIV-1 probe was radiolabeled and incubated with the proteins under study for 20 min at room temperature in binding buffer (10 mM Tris-HCl (pH 7.5), 50 mM NaCl, 10 % (v/v) glycerol, 1 % (v/v) NP-40, 1 mM EDTA, and 0.1 mg/mL poly-dIdC). Samples were run in TGE buffer (24.8 mM Tris base, 190 mM glycine, and 1 mM EDTA) at 200 V for 1 h, and the gel was dried. His-hRelA full-length purified from Sf9 baculovirus cells was used as positive control. Nuclear protein extracts were obtained as described earlier(30) and quantified by Bradford reagent (Bio-rad) to determine the total amount of protein. For supershift reactions, nuclear protein extracts and the anti-RelA antibody were incubated simultaneously for 20 min in the binding buffer in presence of the probe under study.

### IKK2 Expression and Purification

Sf9 insect cells were grown in suspension cultures to a density of 1.5 to 2 × 10^6^ cells/mL prior to infection with baculovirus encoding IKK2 11-669 EE for 48–72 h. Cells were harvested in ice and suspended in buffer (25 mM Tris pH 8.0, 200 mM NaCl, 10 mM imidazole, 10 % glycerol, 5 mM β-mercaptoethanol, and SIGMA protease inhibitor cocktail), treated with 25 μM PMSF immediately before lysis by sonication. The lysate was clarified by centrifugation at 14,000 rpm for 45 min at 4 °C. Supernatant was applied to nickel beads (Ni-NTA resin from BioBharati Life Science, India) equilibrated with lysis buffer. and incubated at 4 °C for 4 h to allow binding before centrifugation at 2000 rpm for 5 min. The protein-bound beads were washed with wash buffer (25 mM Tris pH 8.0, 200 mM NaCl, 30 mM Imidazole, 10 % Glycerol, and 5 mM β-mercaptoethanol) until the washes indicate a protein concentration of less than 0.1 mg/mL by Bradford assay (Bio-rad). The protein was eluted with elution buffer (25 mM Tris pH 8.0, 200 mM NaCl, 250 mM Imidazole, 10 % Glycerol, and 5 mM β-mercaptoethanol) in volumes 1/1000 of the culture size). IKK2 protein was assayed by Bradford assay and in SDS-PAGE, and appropriate fractions were pooled and treated with TEV protease to cleave the His-tag. This protein was incubated with 1 mM ATP in 20 mM MgCl_2_, 20 mM MgCl_2_, 20 mM β-glycerophosphate, 10 mM NaF, and 1 mM sodium orthovanadate for 1 h and further purified in a Superose 6 size-exclusion column equilibrated with 25 mM Tris-HCl (pH 8.0), 250 mM NaCl, 2 mM DTT, and 5 % glycerol. Peak fractions were pooled, concentrated, flash frozen in small aliquots with liquid nitrogen, and stored at -80 °C.

### Hydrogen-deuterium exchange mass spectrometry

Purified IKK2 11-669 EE was incubated with indicated inhibitors for a minimum time-period of 30 minutes in ice to generate the samples for HDX-MS. The protein concentration was 5 μM for ID runs and 10 μM for runs with inhibitor.

HDX-MS was performed using a Waters Synapt G2Si system with HDX technology (Waters Corporation(31). Deuterium exchange reactions were prepared using a Leap HDX PAL autosampler (Leap technologies, Carrboro, NC). D_2_O buffer was prepared by lyophilizing 25 mM Tris pH 8.0, 200 mM NaCl, 1 mM DTT, 0.5 mM EDTA, before being resuspended in 10 mL 99.96% D_2_O immediately before use. Deuterium exchange was measured in triplicate at each time-point (0 min, 30 sec, 1 min, 2 min, and 5 min). For each deuteration time point, 5 μL of protein sample was kept at 25 °C for 5 min prior to addition of 55 μL of D_2_O buffer. The deuterium exchange was quenched by combining 50 μL of the deuteration reaction with 50 μL of ice cold 250 mM TCEP pH 2.5 for 1 min at 1 °C. The quenched sample was injected into a 50 μL sample loop, followed by digestion on an in-line pepsin column (immobilized pepsin, Pierce, Inc.) at 15 °C. The resulting peptides were captured on a BEH C18 Vanguard pre-column, and separated by analytical chromatography (Acquity UPLC BEH C18, 1.7 μM, 1.0 × 50 mm, Waters Corporation) using a 7-85 % acetonitrile gradient in 0.1 % formic acid over 7.5 min, and electrosprayed into the Waters Synapt G2Si quadrupole time-of-flight mass spectrometer. The mass spectrometer was set to collect data in the Mobility, ESI^+^ mode; mass acquisition range 200-2,000 (m/z); scan time 0.4 s. Continuous lock mass correction was accomplished with infusion of leu-enkephalin every 30 s (mass accuracy of 1 ppm for calibration standard). For peptide identification, the mass spectrometer was set to collect data in MS^E^, mobility ESI+ mode instead. Peptides masses were identified from triplicated analyses of 10 μM IKK2 11-669 EE, and data were analyzed using PLGS 2.5 (Waters Corporation). Identification of peptides larger than 1500 Da was done using a minimum number of 250 ion counts for low energy peptides and 50 ion counts for their fragment ions.

The peptides identified in PLGS were analyzed in DynamX 3.0 (Waters Corporation) to determine deuterium uptake. Additional filters in DynamX 3.0 included a cut-off score of 7, minimum products per amino acid of 0.2, maximum MH+ error tolerance of 5 ppm, retention time standard deviation of 5 %, and requiring that the peptide be present in at least two of the three peptide identification runs. The deuterium uptake for each peptide was calculated by comparing the centroids of the mass envelopes of the deuterated samples to the undeuterated controls. For all HDX-MS data, at least 2 biological replicates were analyzed, each with 3 technical replicates. Data are represented as mean values +/-SEM of 3 technical replicates due to processing software limitations, although biological replicates were highly reproducible due to use of the LEAP robot for all experiments. The deuterium uptake was corrected for back-exchange using a global back exchange correction factor (typically 25 %) determined from the average percent exchange measured in disordered termini of various proteins. ANOVA analyses and t tests with a p value cutoff of 0.05 implemented in the program, DECA, were used to determine the significance of differences between HDX data points(32). The peptides reported on the coverage maps are those from which deuterium uptake data were obtained. Deuterium uptake plots were generated in DECA (github.com/komiveslab/DECA) and the data are fitted with an exponential curve for ease of viewing. Data were plotted in DECA as number of deuterons incorporated vs. time (min). The Y-axis limit for each plot reflects the total possible number of amides within the peptide that can exchange. Each plot includes the peptide MH+ value, sequence, and sequential residue numbering.

### Gel-extraction and Mass Spectrometry

IKK2 was incubated with inhibitors for 30 minutes on ice before being separated by 10 % SDS PAGE. The IKK2 bands of the size of 1mm cube were excised from the stained gel and destained by washing initially with 100 μl of 100 mM ammonium bicarbonate for 15 minutes thrice, followed by 100 μl of acetonitrile (ACN) for 15 minutes. The samples were dried in a speedvac, reduced by mixing with 200 μl of 100 mM ammonium bicarbonate-10 mM DTT, and incubated at 56 °C for 30 minutes. The liquid was removed and 200 μl of 100 mM ammonium bicarbonate-55 mM iodoacetamide was added to gel pieces and incubated at room temperature in the dark for 20 minutes. After the removal of the supernatant and one wash with 100 mM ammonium bicarbonate for 15 minutes, same volume of ACN was added to dehydrate the gel pieces. The solution was then discarded and samples were dried in a speedvac. For digestion, enough solution of ice-cold trypsin (0.01 μg/ul) in 50 mM ammonium bicarbonate was added to cover the gel pieces and set on ice for 30 min. After complete rehydration, the excess trypsin solution was removed, replaced with fresh 50 mM ammonium bicarbonate, and left overnight at 37 °C. The peptides were extracted twice by the addition of 50 μl of 0.2 % formic acid and 5 % ACN and mixed with vortex at room temperature for 30 min. The supernatant was removed and saved. A total of 50 μl of 50 % ACN-0.2 % formic acid was added to the sample, which was vortexed again at room temperature for 30 min. The supernatant was removed and combined with the supernatant from the first extraction. The combined extractions are analyzed directly by liquid chromatography (LC) in combination with tandem mass spectroscopy (MS/MS) using electrospray ionization.

### LC-MS-MS

Trypsin-digested peptides were analyzed by ultra-high pressure liquid chromatography (UPLC) coupled with tandem mass spectroscopy (LC-MS/MS) using nano-spray ionization (33). The nanospray ionization experiments were performed using a Orbitrap fusion Lumos hybrid mass spectrometer (Thermo) interfaced with nano-scale reversed-phase UPLC (Thermo Dionex UltiMate™3000 RSLC nano System) using a 25 cm, 75-micron ID glass capillary packed with 1.7-μm C18 (130) BEH™ beads (Waters corporation). Peptides were eluted from the C18 column into the mass spectrometer using a linear gradient (5–80 %) of ACN at a flow rate of 375 μl/min for 1h. The buffers used to create the ACN gradient were: Buffer A (98 % H_2_O, 2 % ACN, 0.1 % formic acid) and Buffer B (100 % ACN, 0.1 % formic acid). Mass spectrometer parameters are as follows: an MS1 survey scan using the orbitrap detector (mass range (m/z): 400-1500 (using quadrupole isolation), 120000 resolution setting, spray voltage of 2200 V, Ion transfer tube temperature of 275 °C, AGC target of 400000, and maximum injection time of 50 ms) was followed by data dependent scans (top speed for most intense ions, with charge state set to only include +2-5 ions, and 5 second exclusion time, while selecting ions with minimal intensities of 50000 at in which the collision event was carried out in the high energy collision cell (HCD Collision Energy of 30%), and the fragment masses where analyzed in the ion trap mass analyzer (With ion trap scan rate of turbo, first mass m/z was 100, AGC Target 5000 and maximum injection time of 35 ms). Protein identification and label free quantification was carried out using Peaks Studio 8.5 (Bioinformatics solutions Inc.)

### IKK2 Kinase Assay

Purified IKK2 was incubated for 30 minutes at room temperature in presence of SC514, TPCA, cpd 65.5 or cpd 65.5.3: The reaction mixture contained 100 ng IKK2, 200 μM cold ATP, 10μCi ^32^P-labeled ATP, 1 μg IκBα; (1-54) and indicated concentration of inhibitors. The samples were quenched directly with SDS PAGE dye, separated on a 12.5 % SDS PAGE gel, exposed to phosphor plates for 24 h, and scanned using STORM860. The gels were also stained afterwards with Coomassie to assess input levels of IKK2 and IκBα.

## ACKNOWLEDGEMENTS

The research was supported by grants NIH CA141722 and American Association for Cancer Research (AACR-18-80-44) grants to GG, UC Tobacco-Related Disease Research Program T29IR0262 to GG and DS. Authors thank Steve Silletti and Majid Ghassemian for help in HDX and MS-MS experiments, and Anup Mazumder for help with Western Blot analyses. We also thank Elizabeth Komives for help with HDX-MS experiments, interpretations of data, and critical reading of the manuscript.

## SUPPLEMENTARY INFORMATION

### SUPPLEMENTARY TABLE

| ChemBridge ID | Ligand efficiency |
| --- | --- |
| 6873991 | -3.59 |
| 4030757 | -2.82 |
| 4030498 | -0.53 |
| 4028668 | -0.48 |
| 4033495 | -0.47 |
| 9117783 | -0.45 |
| 9071338 | -0.42 |
| 4029496 | -0.41 |
| 7768033 | -0.41 |
| 9071985 | -0.38 |
| 4028062 | -0.34 |
| 9145079 | -0.33 |
| 4014357 | -0.33 |
| 4032077 | -0.32 |
| 4028583 | -0.32 |
| 4027040 | -0.30 |
| 4014314 | -0.30 |
| 4026978 | -0.30 |
| 4029463 | -0.30 |
| 6974012 | -0.29 |
| 4013058 | -0.29 |
| 4031557 | -0.29 |
| 9192288 | -0.29 |
| 4027697 | -0.29 |
| 4026988 | -0.28 |
| 4032084 | -0.28 |
| 4004175 | -0.28 |
| 5314018 | -0.26 |
| 4032547 | -0.26 |
| 4014326 | -0.25 |
| 4031471 | -0.25 |
| 6430743 | -0.24 |
| 6626754 | -0.23 |
| 7999350 | -0.22 |
| 5309277 | -0.22 |
| 7656287 | -0.22 |
| 4027609 | -0.22 |
| 7790626 | -0.21 |
| 9034763 | -0.21 |
| 9039007 | -0.21 |
| 9127423 | -0.21 |
| 7784463 | -0.20 |
| 9134173 | -0.20 |
| 9039578 | -0.19 |
| 5489470 | -0.17 |
| 5569363 | -0.17 |
| 5655804 | -0.16 |
| 9001851 | -0.15 |
| 5827311 | -0.15 |
| 5663269 | -0.12 |
| 9193329 | -0.26 |
| 9192288 | -0.23 |
| 9194000 | -0.23 |
| 9195756 | -0.28 |
| 9195757 | -0.27 |
| 9238820 | -0.23 |
| 9241338 | -0.22 |
| 9259528 | -0.21 |
| 9267334 | -0.25 |
| 9276425 | -0.26 |
| 9277493 | -0.35 |
| 9285349 | -0.27 |
| 5649207 | -0.15 |
| 5221976 | -0.15 |
| 5940801 | -0.24 |
| 5192841 | -0.31 |
| 5115429 | -0.22 |
| 5102519 | -0.14 |
| 5107864 | -0.20 |
| 5113658 | -0.18 |
| 5130049 | -0.17 |
| 5137791 | -0.15 |
| 5151224 | -0.14 |
| 5156540 | -0.17 |
| 5194213 | -0.18 |
| 5377737 | -0.21 |
| 5402319 | -0.25 |
| 5403250 | -0.24 |
| 5761623 | -0.16 |
| 7277032 | -0.15 |
| 7659900 | -0.26 |
| 7933991 | -0.20 |
| 9119774 | -0.16 |
| 9142493 | -0.14 |
| 9256557 | -0.22 |
| 5132366 | -0.16 |
| 5226954 | -0.15 |
| 5227793 | -0.18 |
| 5255886 | -0.12 |
| 5355866 | -0.12 |
| 5492396 | -0.21 |
| 5883798 | -0.18 |
| 5963879 | -0.20 |
| 6633931 | -0.12 |
| 6633951 | -0.15 |
| 6636320 | -0.14 |
| 6862662 | -0.22 |
| 7003787 | -0.17 |
| 7015217 | -0.14 |
| 7600068 | -0.14 |

