## Supplementary Figures for "Discovery of an IKK2 Site that Allosterically Controls Its Activation"

### Supplementary Figure 1.

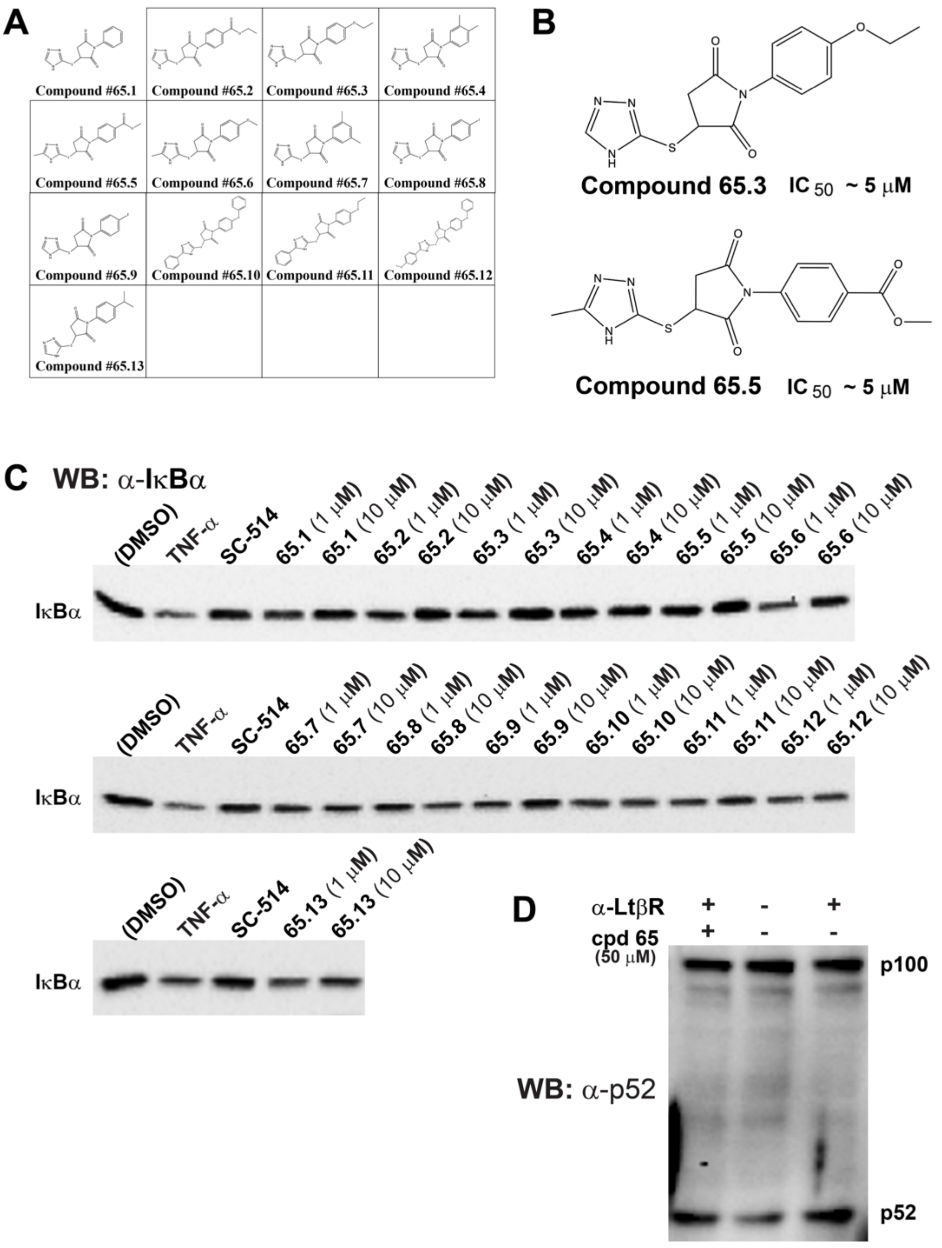


**Supplementary Figure 1**.
Expanded structure-activity relationship analysis via second-generation derivatives of cpd 65. Effect of 13 commercially available compounds (cpds 65.1 to 65.13) (**A**) on IκBα level in TNF-α treated cells. Two compounds (cpd 65.3 and cpd 65.5) (**B**) showed significant reduction (**C**) in IκBα degradation. (**D**) Cpd 65.5 treatment showed no discernible effect on p100 processing in MEF cells.

**Supplementary Figure 2A.**

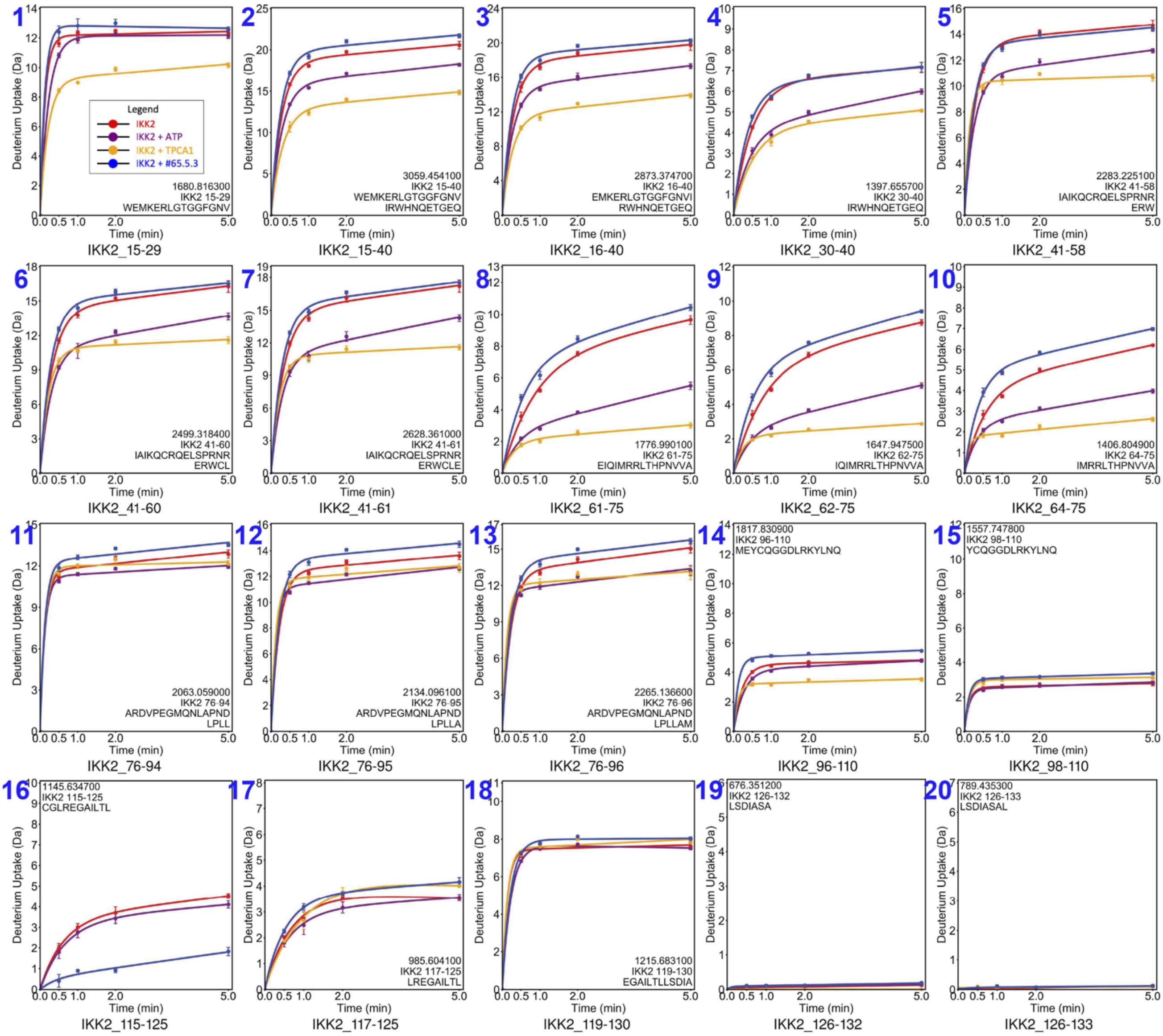


**Supplementary Figure 2A.** Panels 1-20.

HDX-MS profiles showing comparison of the protection and deprotection of H-D exchange in IKK2 (residue ranges ~ 15-133) in its free (red), partnered with ATP (purple), ATP-competitive inhibitor TPCA (orange), and cpd 65.5.3 (blue) states.

**Supplementary Figure 2B.**

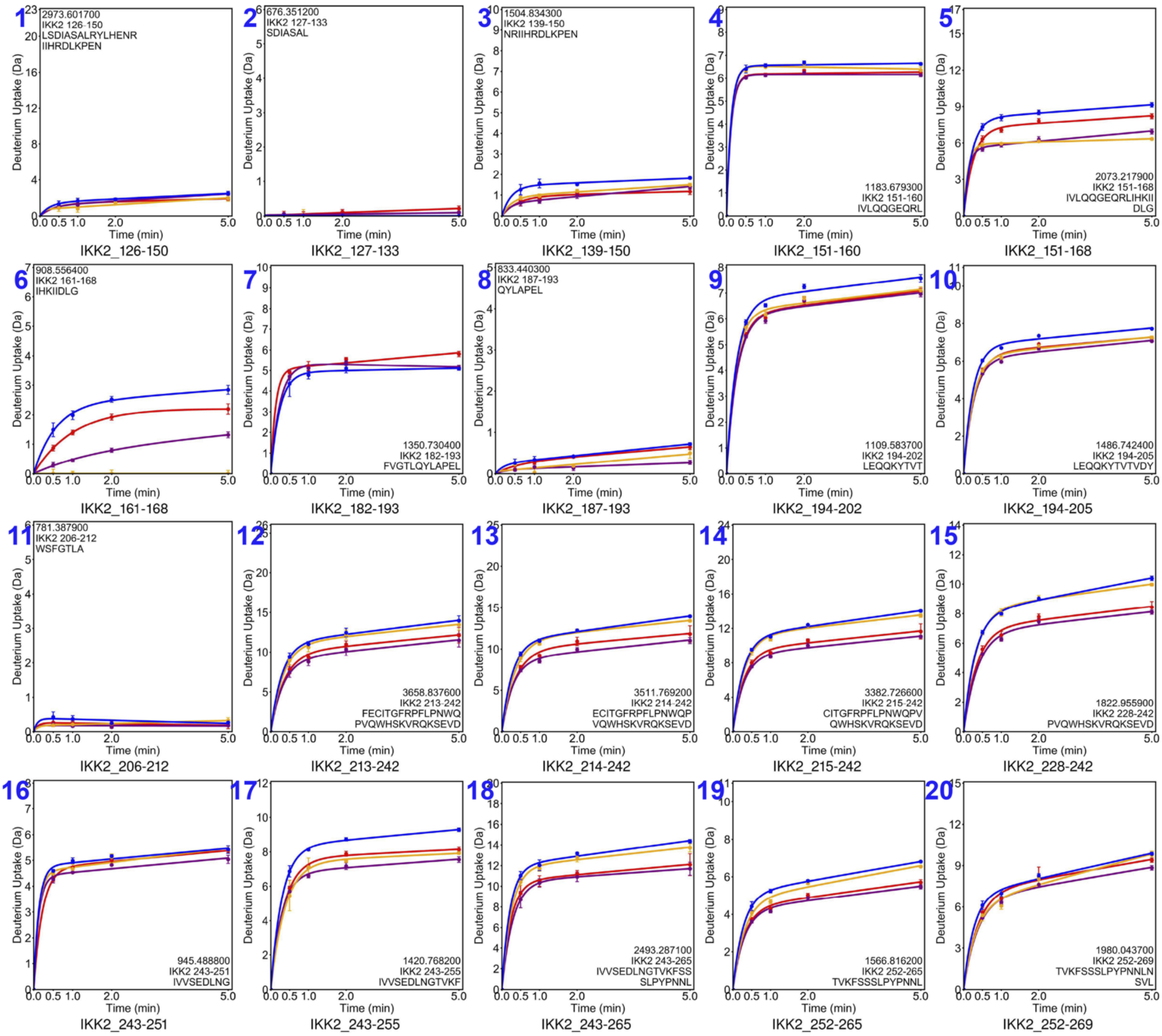


**Supplementary Figure 2B.** Panels 1-20.

HDX-MS profiles showing comparison of the protection and deprotection of H-D exchange in IKK2 (residue ranges ~ 126-269) in its free (red), partnered with ATP (purple), ATP-competitive inhibitor TPCA (orange), and cpd 65.5.3 (blue) states.

**Supplementary Figure 2C.**

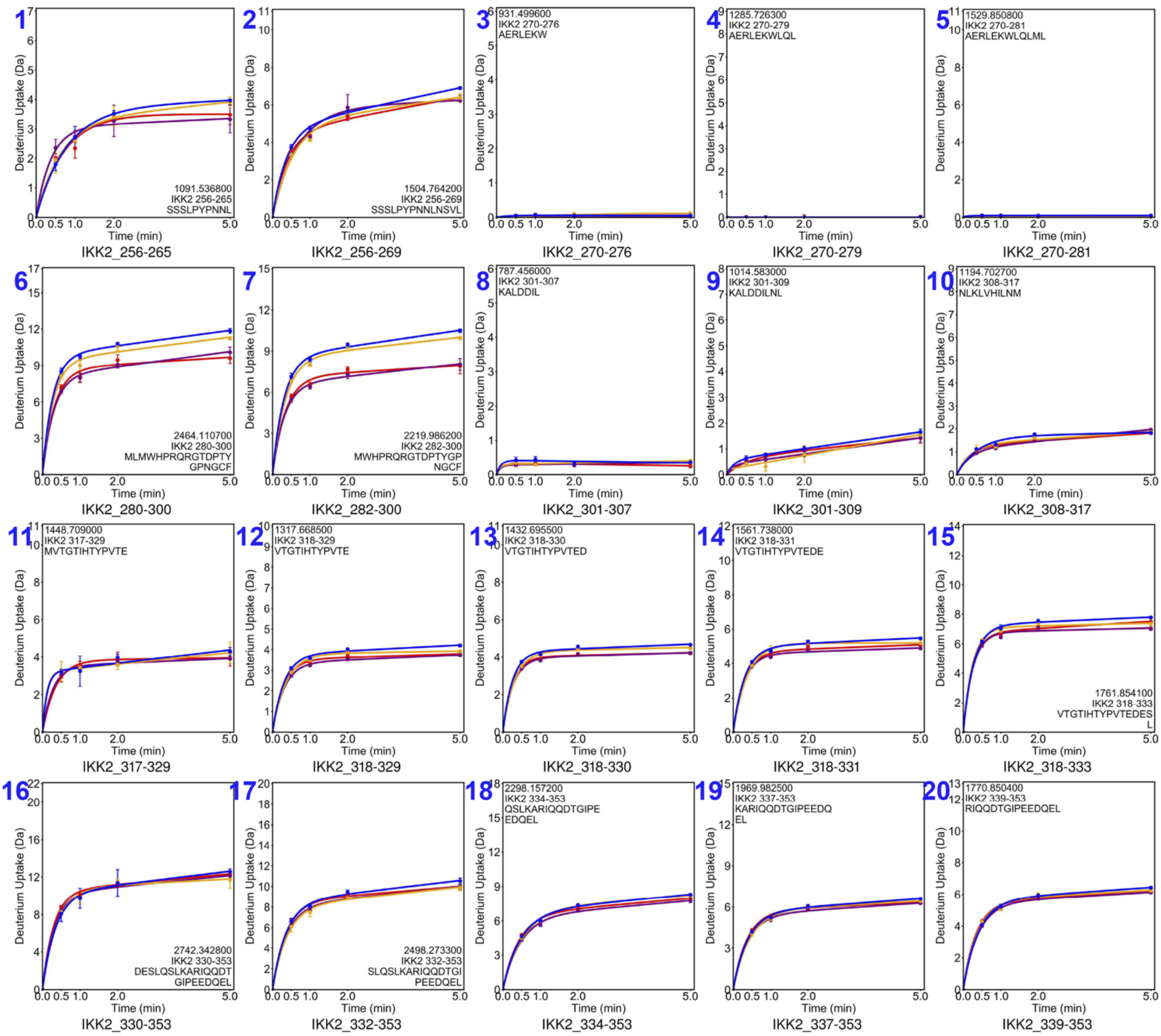


**Supplementary Figure 2C.** Panels 1-20.
HDX-MS profiles showing comparison of the protection and deprotection of H-D exchange in IKK2 (residue ranges ~ 256-353) in its free (red), partnered with ATP (purple), ATP-competitive inhibitor TPCA (orange), and cpd 65.5.3 (blue) states.

**Supplementary Figure 2D.**

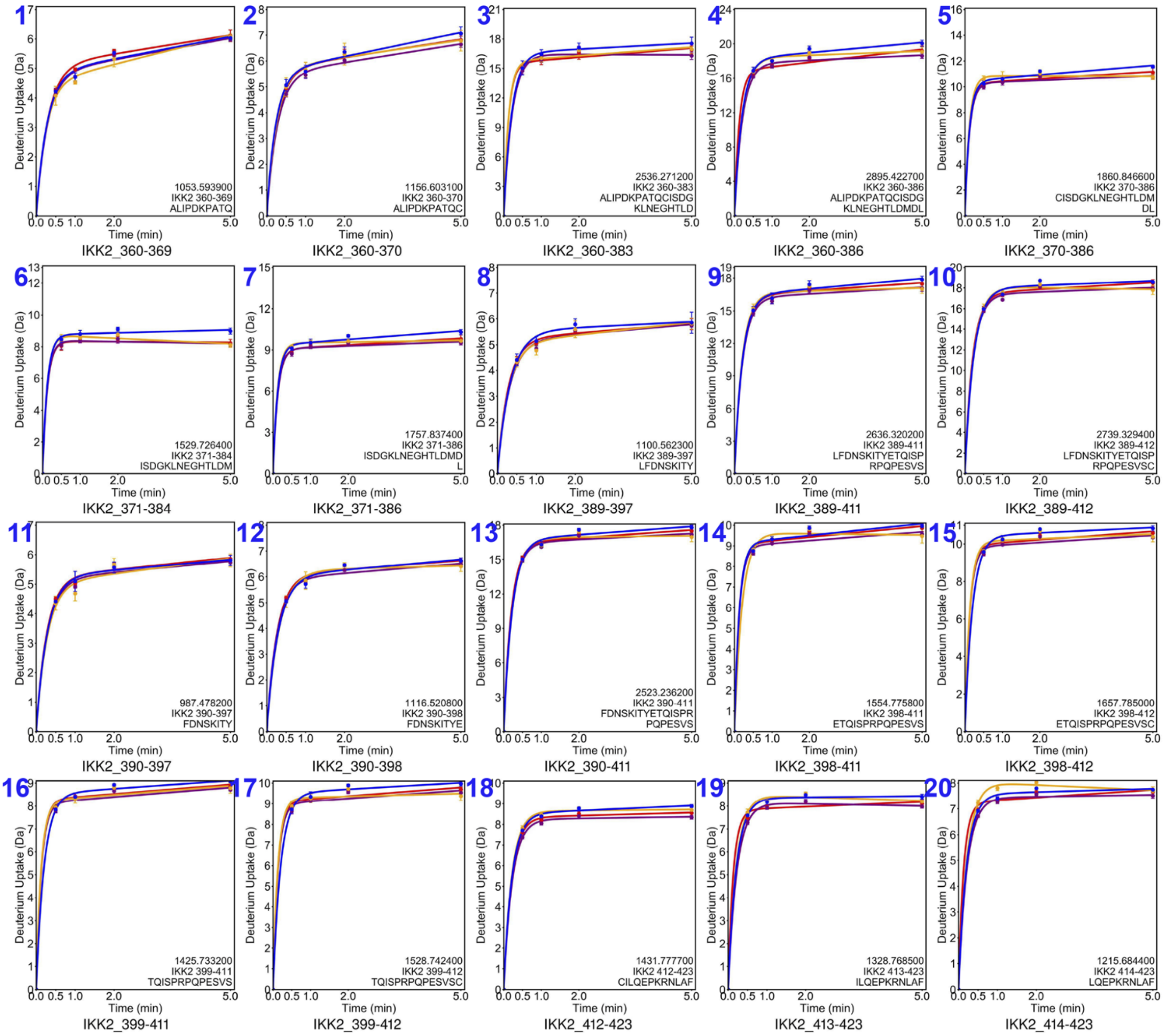


**Supplementary Figure 2D.** Panels 1-20.

HDX-MS profiles showing comparison of the protection and deprotection of H-D exchange in IKK2 (residue ranges ~ 360-423) in its free (red), partnered with ATP (purple), ATP-competitive inhibitor TPCA (orange), and cpd 65.5.3 (blue) states.

**Supplementary Figure 2E.**

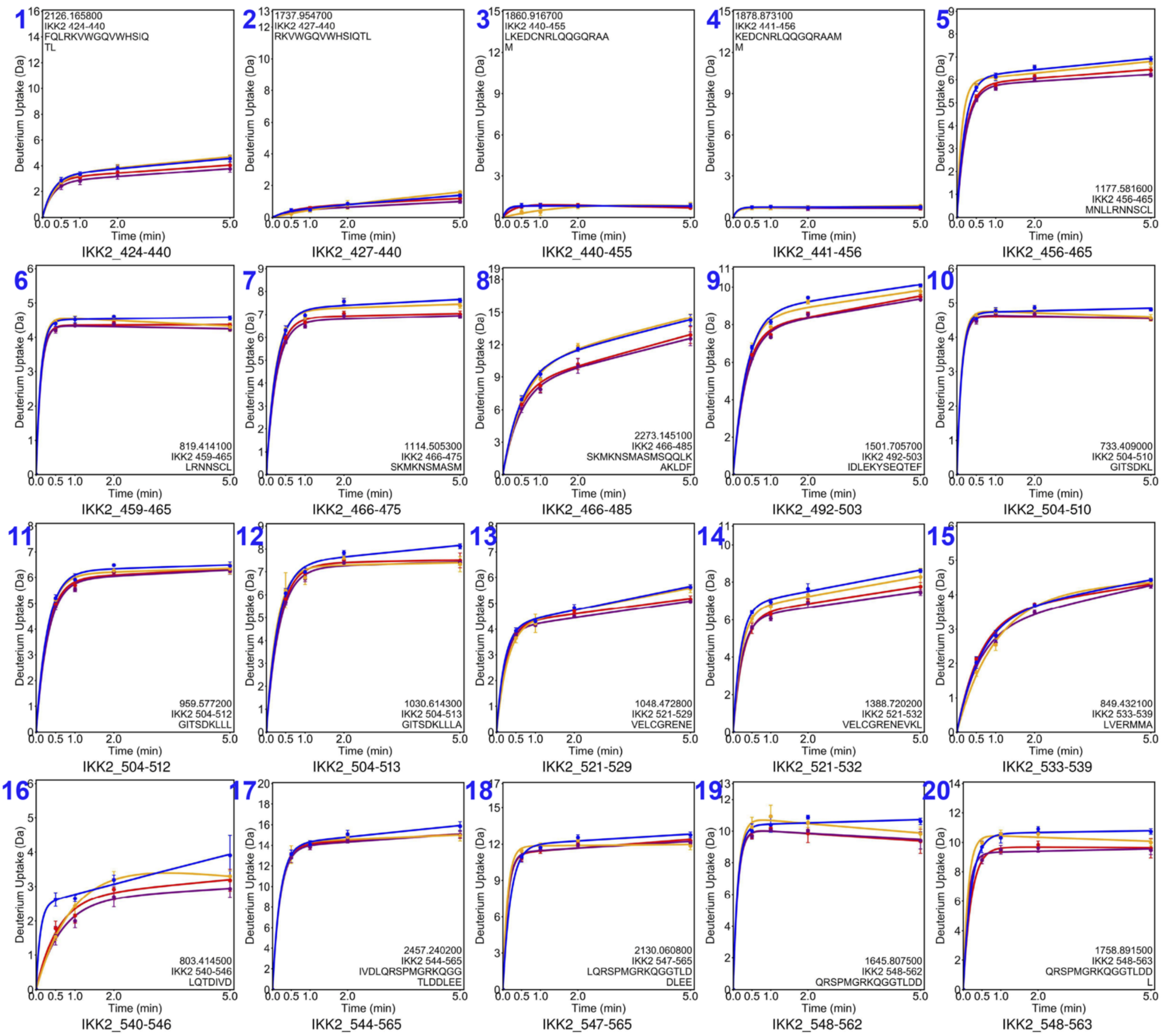


**Supplementary Figure 2E.** Panels 1-20.
HDX-MS profiles showing comparison of the protection and deprotection of H-D exchange in IKK2 (residue ranges ~ 424-563) in its free (red), partnered with ATP (purple), ATP-competitive inhibitor TPCA (orange), and cpd 65.5.3 (blue) states.

**Supplementary Figure 2F.**

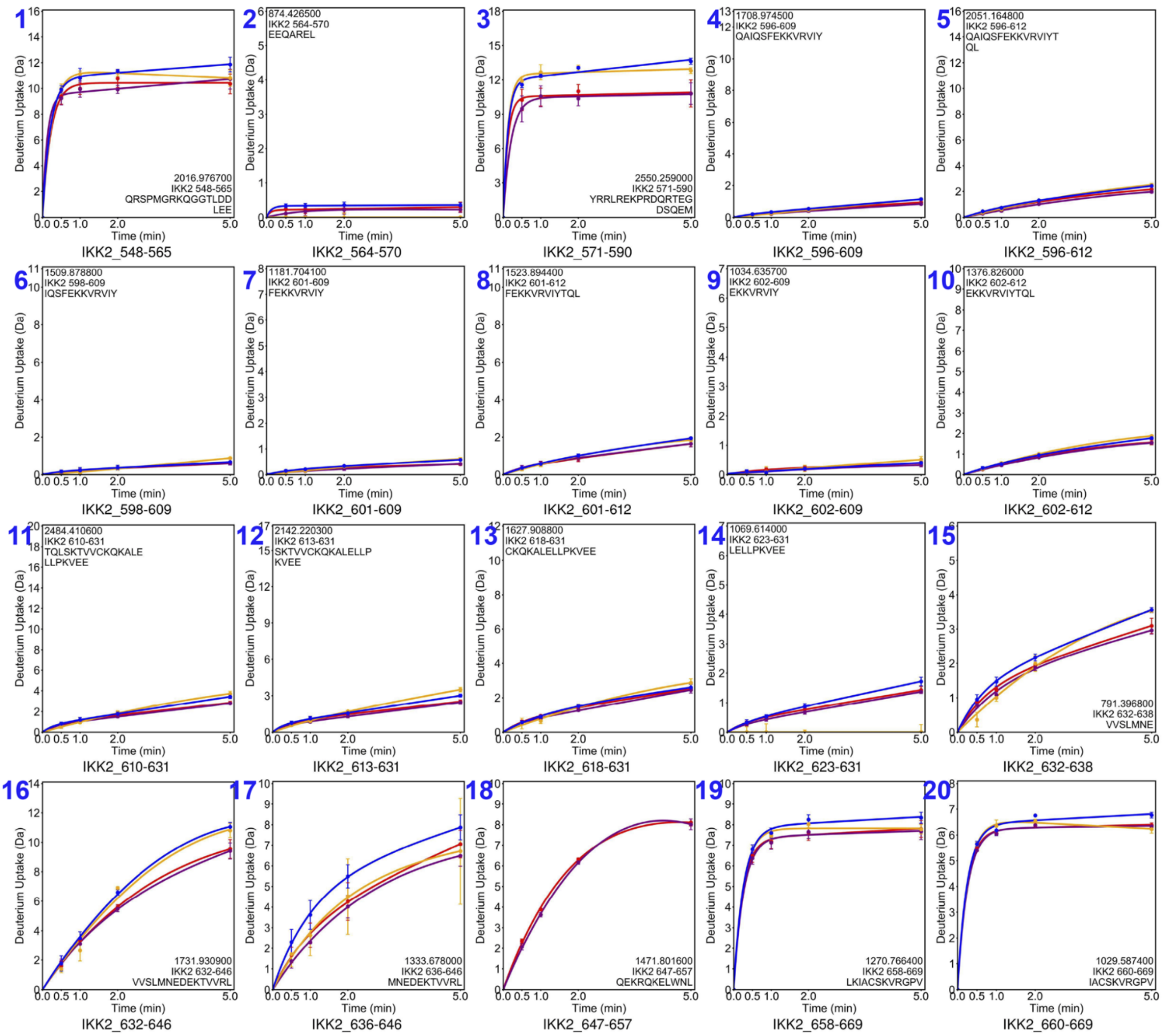


**Supplementary Figure 2F.** Panels 1-20.
HDX-MS profiles showing comparison of the protection and deprotection of H-D exchange in IKK2 (residue ranges ~ 548-669) in its free (red), partnered with ATP (purple), ATP-competitive inhibitor TPCA (orange), and cpd 65.5.3 (blue) states.

**Supplementary Figure 3.**

**A**

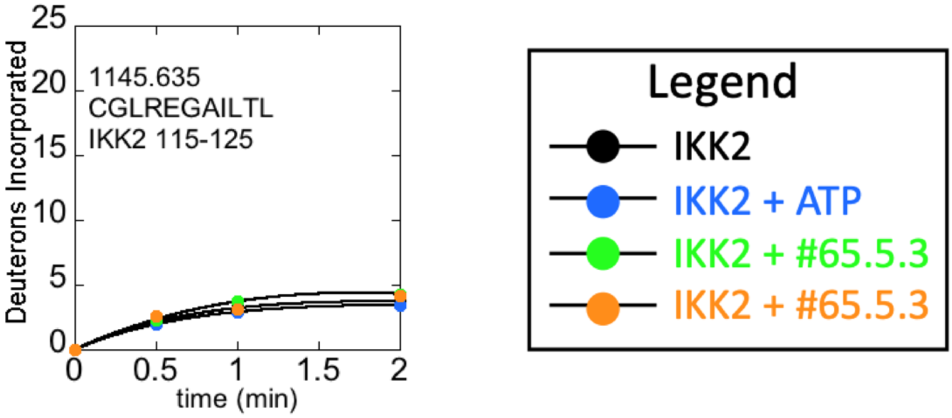


**B**

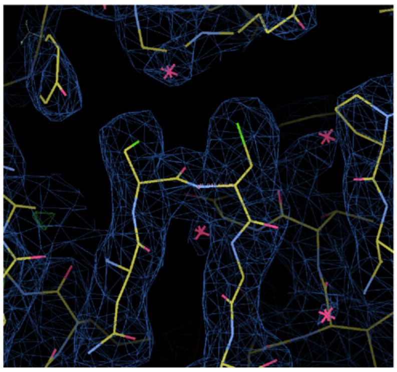

**C**

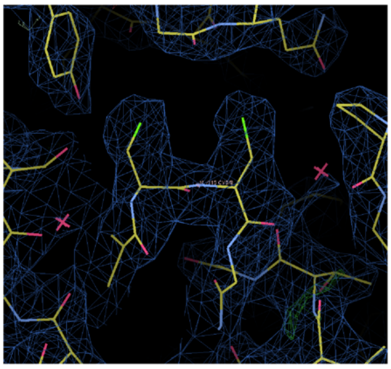

**Supplementary Figure 3.**
(**A**) H-D exchange shows no deprotection in IKK2 peptide 115 to 125 by ATP or inhibitor 65.5.3 in absence of DTT. This peptide includes the cysteine 115 of the di-Cys 114/115 motif. Comparison of behavior of fragment 117-125 without cysteine to 115-125 (**B**) Density map showing the position of cysteine 114 and 115 sidechains in inactive monomer of IKK2 in pdb model 4kik. (**C**) Density map showing the position of cysteine 114 and 115 sidechains in active monomer of IKK2 in pdb model 4kik.

**Supplementary Figure 4.**

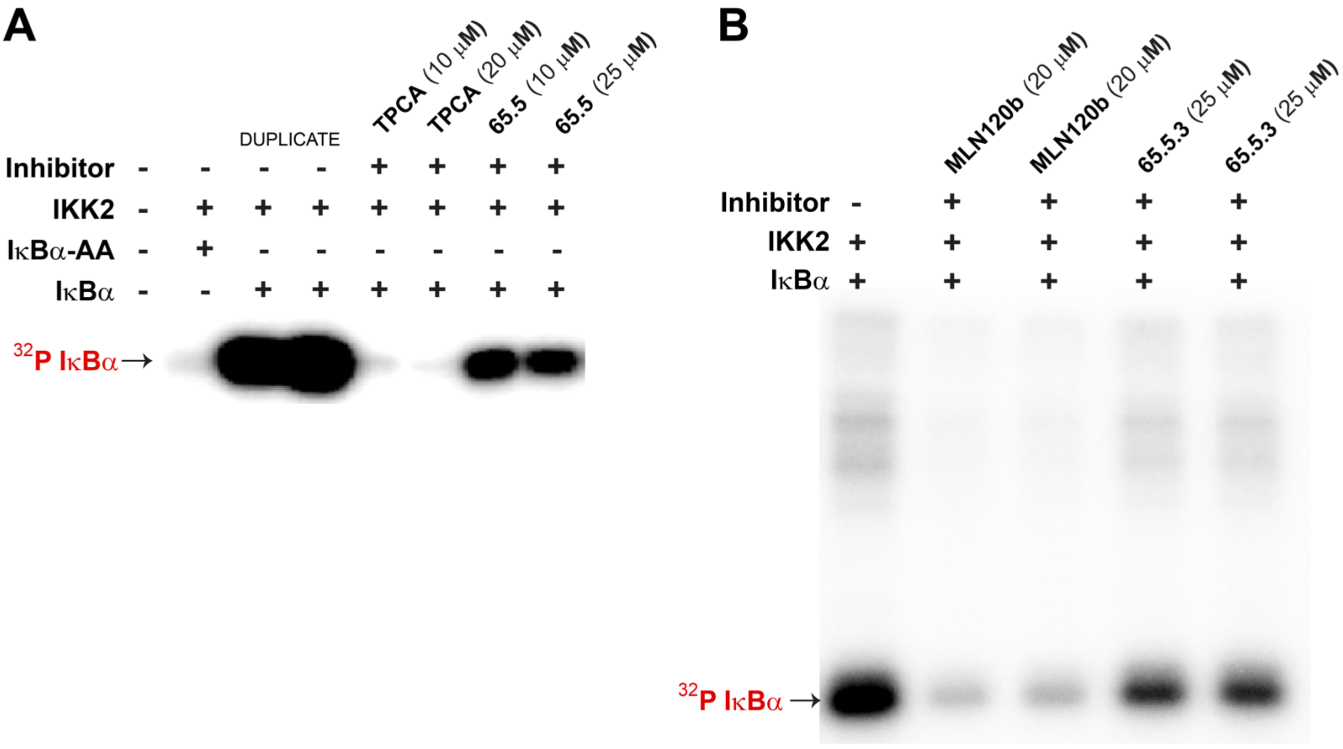


**Supplementary Figure 4.** Inhibitory effects of cpds 65.5 and 65.5.3 on IKK2. Reaction mixtures (100 ng IKK2, 200 μM ATP, 10 μCi $^{32}$P-labeled ATP, 1 μg IκBα) were incubated with TPCA and cpd 65.5 (**A**), and MLN120b and cpd 65.5.3 (**B**) for 30 minutes at RT, quenched with SDS-PAGE dye, separated in SDS-PAGE, and autoradiographed. Phosphorylation of IκBα by IKK2 is nearly abolished by TPCA or MLN120b (ATP-competitive inhibitors) and significantly reduced with cpds 65.5 (panel A) and 65.5.3 (panel B).
